# Comprehensive Phenotyping of Extracellular Vesicles in Blood of Healthy Humans – Insights into Cellular Origin and Biological Variability

**DOI:** 10.1101/2024.07.04.602156

**Authors:** Marija Holcar, Ivica Marić, Tobias Tertel, Katja Goričar, Urška Čegovnik Primožič, Darko Černe, Bernd Giebel, Metka Lenassi

## Abstract

Despite immense interest in biomarker applications of extracellular vesicles (EVs) from blood, our understanding of their physiological population in healthy humans remains limited. Using imaging and multiplex bead-based flow cytometry, we comprehensively quantified circulating EVs with respect to their cellular origin in a large cohort of healthy blood donors. We assessed coefficients of variations to characterise their biological variability and explored demographic, clinical, and lifestyle factors contributing to this variability. Cell-specific circulating EV subsets show a wide range of concentrations, which do not directly reflect concentrations of blood cells, indicating diverse patterns of EV subset release and/or uptake, even for EVs originating from the same cell type. Interestingly, tetraspanin+ circulating EVs largely originate from platelets and to a lesser extent from lymphocytes. PCA and association analyses demonstrate high biological inter-individual variability in circulating EVs across healthy humans, which can be only partly explained by the influence of sex, menopausal status, age and smoking on specific circulating EV and/or tetraspanin+ circulating EV subsets. No global influence of the explored subject’s factors on circulating EVs was detected. Our findings provide the first comprehensive, quantitative data towards the cell-origin atlas of blood EVs, with important implications in the clinical use of EVs as biomarkers of disease.

## Introduction

Plasma, the liquid part of the blood, is the predominant source for extracellular vesicle (EVs) biomarker, therapeutic and functional studies (Royo et al. 2020; Beetler et al. 2023). Despite huge interest, we have limited knowledge of the circulating EV population in healthy humans. Their total concentration was estimated from 10^6^ EVs/mL with flow cytometry to up to 10^13^ EVs/mL with nanoparticle tracking analysis (NTA) (Johnsen et al. 2019). With respect to the cellular source of circulating EVs, platelet-derived and erythrocyte-derived EVs are in the range of 10^7^ EVs/mL, while quantification of EVs derived from leukocyte subsets in plasma is mostly missing (Arraud et al. 2014, Bettin et al. 2022, Kumar et al. 2024). Digital quantification based on the long RNA profiles estimated that 99.8% of circulating EVs originate from hematopoietic cells, mostly from platelets, B cells, and CD4 T cells (Li et al. 2020), but the extent to which RNA is packed into more numerous small EVs is controversial (Tosar, Witwer, and Cayota 2021; Mosbach et al. 2021). Several recent proteomics studies have provided comprehensive protein composition of circulating EVs in healthy humans (Rai et al. 2024; Vallejo et al. 2023; Muraoka et al. 2022), but there is a lack of quantification at the single vesicle level. Thus there is a key need for comprehensive single vesicle quantification of circulating EV population in healthy humans.

Importantly, the relationship between concentrations of specific blood cells and circulating EVs is unclear. At any time, the steady-state concentration of circulating EVs is the equilibrium of EV secretion and uptake, but direct measurements in human blood are missing. To address this, Auber and Svenningsen integrated available data on circulating EVs and blood cells and estimated the highest EV secretion rates for monocytes and the lowest for erythrocytes, with platelets in the middle (Auber and Svenningsen 2022). EV uptake studies in animal models identified macrophages and endothelial cells in mice, and B cells in macaques, as largely responsible for EV clearance from blood in a few minutes to a few hours (Driedonks et al. 2022; Eitan et al. 2017; Loconte et al. 2023; Matsumoto et al. 2021). A more direct approach to studying the relationship between specific blood cell concentrations and levels of circulating EVs, with respect to their cell of origin, is to perform quantification and association studies on a large healthy human cohort.

Another fairly neglected aspect of circulating EVs is intrinsic biological variation across healthy humans. Understanding biological variation is fundamental for circulating EVs’ clinical application, especially for interpretation of EV-based disease biomarker readouts. The biological variation of commonly examined blood plasma quantities is widely documented, most showing higher inter-individual than intra-individual variation (Aziz et al. 2019; Widjaja et al. 1999). To improve clinical decisions, reference intervals for most haematological quantities are also stratified by sex and/or age (L. Van Pelt et al. 2022; Ozarda, Higgins, and Adeli 2019). In the case of circulating EVs, current studies imply the influence of age, sex, smoking and physical activity. Individual studies on small circulating EVs in healthy humans showed decreased blood extracellular particle concentration with advancing age (Eitan et al. 2017), increased levels of tetraspanin+ circulating EVs during cycling exercise (Brahmer et al. 2019) and changes in tetraspanin+ circulating EVs protein profiles in smokers, depending on sex (Bæk, Varming, and Jørgensen 2016). Flow cytometric studies of large circulating EVs showed higher levels of EVs derived from platelets and endothelial cells in females compared to males (Gustafson et al. 2015), and the influence of age, sex and smoking status on specific EV subsets (Enjeti et al. 2017). So far, there are no comprehensive studies of circulating EVs biological variation in large healthy human cohorts, and the impact of demographic, clinical, and lifestyle factors on the latter.

The aim of this study was to improve the understanding of physiological circulating EVs by comprehensively quantifying circulating EVs with respect to their cell of origin in healthy humans, exploring the biological variability and demographic, clinical, and lifestyle sources of the observed variability. To this end, we performed surface protein phenotyping of EVs in blood plasma and in isolated tetraspanin+ circulating EVs in a large well-characterised cohort of blood donors, using imaging and multiplex bead-based flow cytometry. We calculated coefficients of variation (CV) for concentrations of circulating EV subsets and marker expression levels of tetraspanin+ circulating EVs to characterise biological variability and conducted association analyses to explore the influence of demographic, clinical, and lifestyle factors on the latter.

## Materials and methods

### Study design and subjects

We performed a cross-sectional study that included 208 adults who donated blood at the Blood Transfusion Centre of Slovenia between July 27 and November 23, 2020. The study design is represented in Figure 1. The study subjects were equally distributed between sexes (M/F) and four age groups (18-29 years; 30-44 years; 45-54 years; 55-64 years). All subjects were Caucasians and had eaten a non-fatty meal prior to blood donation according to the general guidelines for blood donation. Only individuals who exhibited no signs of illness according to the Slovenian blood donation eligibility requirements, which are aligned with The European Directorate for the Quality of Medicines & HealthCare guidelines (EDQM 2023), and were not taking any medications or hormonal contraception at the time, were included in the study. The eligibility of study subjects was further determined through a comprehensive medical interview and examination always conducted by the same physician. Collected fully anonymised data included the subjects’ demographics (age, sex), clinical (body temperature, height, weight, body mass index (BMI), blood pressure, blood pulse, menopause status), and lifestyle (exercise regimen, time since last exercise, smoking status) factors. Written informed consent was obtained from all patients. The study was approved by the Slovenian National Ethics Committee (0120-209/2020/3) and conducted in accordance with the Declaration of Helsinki.

**Figure 1:**
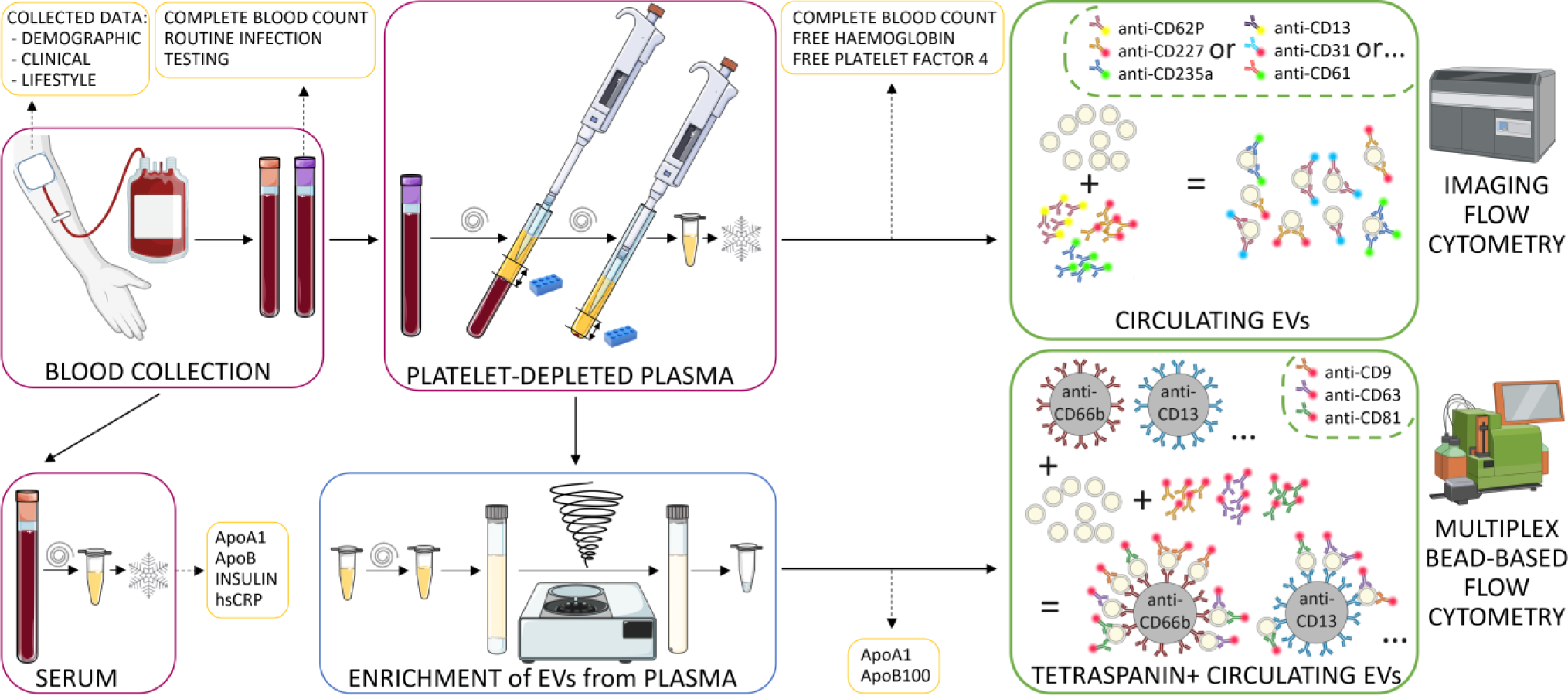
Schematic representation of the study workflow. ApoA1 – apolipoprotein A1; ApoB – apolipoprotein B; hsCRP – highly sensitive measurement of C-reactive protein. Image created by Biorender.com and Smart.servier.com.

### Blood collection and routine laboratory analysis

Blood collection was performed during routine blood donation from a satellite bag into serum (5 mL, BD Vacutainer, UK) and plasma (anticoagulant ethylenediaminetetraacetic acid; EDTA K3/7.5 mL, S-Monovette, Sarstedt, Germany) vacutainer tubes. All blood donations occurred between 7-11 a.m. from a cubital vein, using a butterfly system with a 16G needle and a tourniquet. Whole blood was then routinely analysed to determine complete blood count and exclude common infections (HIV, HBV, HCV, and syphilis).

### Platelet-depleted plasma and serum preparation

Serum was prepared by leaving whole blood to coagulate for 30 min at room temperature, followed by centrifugation for 10 min at 2000 x g (Eppendorf Centrifuge 5702, Hamburg, Germany). Serum was then aliquoted into 1 mL portions and frozen at -80°C until further analysis. Plasma was prepared according to the guidelines of the International Society for Thrombosis and Haemostasis protocol for generating platelet-depleted plasma (Lacroix et al. 2012; Nieuwland and Siljander 2024) as follows: after collection, blood was mixed with an anticoagulant by gently turning the tube six times and then placed in an upright position in the tube rack. Next, two 15-minute consecutive centrifugations at room temperature and 2500 × g were performed without a brake (Eppendorf Centrifuge 5804, Hamburg, Germany). After the first centrifugation, plasma was transferred into a fresh tube. Special care was taken to leave 1 cm of plasma above the sedimented cells after both centrifugation steps. Complete blood count measurements were repeated in plasma (Cell-Dyn Ruby Hematology Analyzer, Abbott, Chicago, IL, USA), then plasma was aliquoted into 1 mL portions and frozen at -80°C until further analysis. Plasma and serum preparations were performed in the same facility as blood donations, always by the same person, and aliquoted and frozen within 4 hours of blood donation. All information regarding plasma and serum preparation is reported also as the MIBlood-EV reports.

### Measurement of lipoproteins, C-reactive protein, and insulin

The lipid profile, highly sensitive measurements of concentrations of C-reactive protein (hsCRP), and insulin were measured in the defrosted serum. The lipid profile was determined by routine nephelometric measurements of apolipoprotein A1 (ApoA1) and apolipoprotein B (ApoB) concentrations using a routine analyser (Atellica NEPH 630, Siemens, Germany) and commercial reagents (N AS APOAI, N AS APOB, Siemens, Germany).

To measure the concentrations of CRP and insulin, we used commercially available ELISA kits: hs-CRP Human C-Reactive Protein ELISA Kit (#RAB0096, Sigma-Aldrich, Germany) and Mercodia Insulin ELISA (#10-1113-01, Mercodia, Sweden). Both ELISAs were performed in duplicates, according to the manufacturer’s instructions.

### Measurement of free haemoglobin and plasma factor 4

Free plasma haemoglobin and platelet factor 4 (PF4) concentrations were measured in defrosted plasma. Free haemoglobin was measured spectrophotometrically by measuring A_380_, A_415_, and A_450_. Concentration was calculated according to the Harboe method with Allen correction: Hb(g/l) = (167.2xA_415_ - 83.6xA_380_ - 83.6xA_450_) x (1/1,000) x dilution in dH_2_0 (Han, Serrano, and Devine 2010). Samples of plasma with a haemoglobin concentration greater than 0.5 g/L were considered haemolysed (Han, Serrano, and Devine 2010). The concentration of PF4 was measured using a commercially available ELISA kit (#EHPF4, Invitrogen, USA) in duplicate, according to the manufacturer’s instructions.

### Imaging flow cytometry analysis

Following defrosting, aliquots of plasma samples were centrifuged at 10,000 x g for 10 min at 4°C to remove larger debris and cryoprecipitates. The supernatant was carefully transferred to new containers to avoid contamination. For antibody staining, aliquots of 10 µL plasma were incubated with fluorescently labelled antibodies targeting EV surface proteins. Up to 4 antibodies were used per measurement and a combination was selected that resulted in a maximum of one double staining on one EV. The cellular origin of chosen markers (Supplementary Table 1) was determined based on established markers recognised in flow cytometry as indicative of specific cells. (“CD Marker Handbook Human and Mouse” 2016). A detailed description of the amounts used and allocation of antibodies to assays is given in Supplementary Tables 2 and 3. The antibodies were diluted in 10 µL PBS and incubated with the plasma aliquots at room temperature for 1 hour to allow for optimal binding. Following incubation, the samples were diluted to a final volume of 100 µL with PBS for analysis (10x dilution). In adherence to the MIFlowCyt-EV criteria for EV analysis (Welsh et al. 2020), we included various controls in our experimental design. These included buffer-only controls, buffer plus antibody controls, fluorochrome-conjugated isotype controls, and detergent lysis controls using 2% NP-40 in PBS. The checklist for MIFlowCyt-EV can be found in the supplement.

All stained samples were analysed using an AMNIS ImageStreamX Mark II Flow Cytometer (AMNIS/Luminex, Seattle, WA, USA) equipped with an autosampler for U-bottom 96-well plates (Corning Falcon, cat 353077) after molecules of equivalent soluble fluorophores (MESF) calibration for fluorescence quantification as described previously (Tertel, Görgens, and Giebel 2020). The results can be found in the Supplementary Figure 1. The cytometer settings were adjusted to an acquisition time of 5 min per well, utilising a 60x magnification and a low flow rate (0.3795 ± 0.0003 μL/min). This setup allowed for high-resolution imaging of EVs without the need for bead removal, as per our previous protocols (Görgens et al. 2019; Tertel et al. 2020). The laser and filter configuration are given in Supplementary Table 4 and the compensation matrix in Supplementary Table 5. Events characterised by low side scatter (<10,000) and a fluorescent intensity higher than 300 were considered for concentration calculations, as detailed in our previous studies (Tertel, Görgens, and Giebel 2020).

### Enrichment of small EVs with sucrose cushion ultracentrifugation (sUC)

EVs were enriched from plasma following our previously established protocol (Holcar et al. 2020). Briefly, upon defrosting 1 mL aliquots of plasma on ice, the samples were centrifuged at 10,000 × g for 20 min at 4°C to remove large extracellular vesicles and any residual cell debris. 900 μL of supernatant was then diluted with 8.5 mL of particle-free Dulbecco’s phosphate-buffered saline (DPBS, Sigma-Aldrich, USA) and carefully layered over 2 mL of 20% sucrose (Merck Millipore, USA) in polypropylene tubes (Beckman Coulter, USA). The samples were ultracentrifuged at 100,000 × g for 135 min at 4°C (MLA-55 rotor, Beckman Coulter, USA). After removal of the supernatant, the pellet containing enriched EVs was resuspended in 60 µL of DPBS, aliquoted into 20 µL samples, and stored at -80 °C. All EV enrichments were performed by the same person in the same laboratory within two months.

### Quantification of EVs by nanoparticle tracking analysis (NTA)

Enriched EVs samples were diluted in DPBS to 1 x 10^7^-10^9^ EVs/mL and examined using a NanoSight NS300 (NanoSight Ltd., UK) equipped with a 488 nm blue laser and automated sample assistant for automated loading of the samples into the instrument. Duplicate samples were pipetted into 96-well plates and five 60 s-long videos were captured for each replicate at camera level 15. After visual inspection, videos with visible artefacts were removed and three videos from each replicate were chosen for analysis. To correct for potential inter-plate variation in NTA measurements, an internal reference EV sample was added in duplicates to each plate and used for the normalisation of analysed EV samples.

Raw laser scattering and particle movement data were analysed using NTA software (version 3.3; NanoSight Ltd., UK). Automatic settings were selected for the maximal track length, minimum expected particle size and blur, minimum track length was set to 10, detection threshold to 5, and sample viscosity to the corresponding viscosity for water and temperature to 25°C. The output data are presented as EV size (modal hydrodynamic diameter in nm) and EV concentration (number of EVs enriched from 1 mL of plasma in particles/mL of plasma). The analytical coefficient of variations (% CV) was calculated from the variations of the combined six measurements of each sample. All analyses were performed by the same person in the same laboratory within eight months after EV enrichment.

### Multiplex bead-based flow cytometry analysis

For semi-quantitative analysis of 37 different surface protein markers, which co-express with at least one of the tetraspanins of enriched circulating EVs, we used a commercial multiplex bead-based flow cytometry kit (MACSPlex Exosome Kit, human, Miltenyi Biotec, Germany) according to the manufacturer’s instructions and analysed the samples with MACSQuant Analyzer 10 (Miltenyi Biotec, Germany). In short, 1 × 10^9^ (according to NTA analysis) of enriched EVs were incubated overnight in the dark at room temperature with a mix of polystyrene capturing beads (each bead coated with one type of specific antibodies) for the detection of 37 EV surface markers and 2 isotype controls. After the incubation, a cocktail of detection antibodies against tetraspanins (mix of anti-CD9, anti-CD63, and anti-CD81 antibodies), all conjugated with APC was added, and mean fluorescence intensity (MFI) was measured (MACSQuant Analyser 10, Miltenyi Biotec, Germany). As a control, MACSPlex Buffer alone was incubated with beads and detection antibodies. The results of MFI measurements were normalised according to the manufacturer’s instructions. First, the background was corrected by subtracting the MFI of the buffer-only sample. Next, the MFIs of each sample were normalised by their respective normalisation factors, calculated by dividing the mean MFI of CD9, CD63, and CD81 signal of the respective sample with the mean MFI of CD9, CD63, and CD81 signal of all samples. When referring to measured MFI levels, we introduced the term ‘marker expression’ level, as MFIs reflect both changes in concentration of specific tetraspanin+ circulating EV subset and changes in expression levels of measured protein markers on those EV subsets.

### Statistical analysis

Statistical analyses were performed using IBM SPSS Statistics, version 27.0 (IBM Corporation, Armonk, NY, USA) and GraphPad Prism, version 9.4.0 (GraphPad Software, La Jolla, CA, USA).

Continuous variables were described as the median and interquartile range (25%–75%), whereas categorical variables were described using frequencies. Non-parametric Kruskal-Wallis and Mann-Whitney tests were used to compare the distribution of continuous variables between different subject groups. Spearman’s rho correlation coefficient (ρ) was used to assess correlations between continuous variables. For significant associations, ρ between 0.3 and 0.6 or -0.3 and -0.6 was considered fair, and above 0.6 or below -0.6 was moderate or strong (Akoglu 2018). All statistical tests were two-sided, and the level of significance was generally set to 0.05, except for analyses of multiple EV subsets or surface markers on tetraspanin+ EVs, where Bonferroni correction was considered. As 25 or 23 EV subsets or EV-surface proteins were evaluated using imaging or multiplex bead-based FC, respectively, P-values below 0.002 were considered statistically significant in this part of the analysis. % CV was calculated as (standard deviation/sample mean) × 100 to measure the relative dispersion of data points in a data series around the mean. We used Principal Component Analysis (PCA) for dimensionality reduction and data visualisation when analysing the influence and co-dependence of 25 EV subset concentrations or 23 EV-surface protein expressions. PC1 and PC2 represent the first and second principal components, respectively, which are linear combinations of the original variables that capture the maximum variance in the data. Loadings refer to the weights assigned to each original variable in the calculation of principal components, indicating their contribution to the variance observed in the data.

## Results

### Quantification of EV populations, with respect to their cell of origin, in the blood of healthy humans

To gain insight into the physiological EV population in the blood of healthy humans, we first recruited 208 adult blood donors who satisfied all general requirements for Slovenian blood donation eligibility (including the absence of any sign of illness) and were not taking any medications or hormonal contraception at the time. They all tested negative for HIV, HBV, HCV, and syphilis infections and had normal CRP levels, measured at high sensitivity, implying an absence of inflammation or infection. Study subjects were further characterised for demographic, clinical, and lifestyle factors by in-depth interviews, examinations and routine blood tests (Table 1). Study subjects matched by sex and age (18-65 years, median age 44.5 years) had a median BMI representative of Slovenian population (25.6 (25-75%: 22.8-28.1) kg/cm^2^) (Jakše, Godnov, and Pinter 2022). Their body temperature, blood pressure and blood pulse were inside respective reference ranges. The study subjects’ metabolism balance was described by serum lipoproteins (ApoA1 and ApoB) and insulin concentrations. No impact of lipoprotein or insulin concentrations on any of the EV characteristics studied here was observed (Supplementary Table 6).

**Table 1:** Study subject’s characteristics.

|  | Unit / Category | Median (25-75%) | N (%) |
| --- | --- | --- | --- |
| Sex | Male |  | 104 (50.0) |
|  | Female |  | 104 (50.0) |
| Age | All subjects / years | 44.5 (29.3-54.8) |  |
|  | 18-29 years |  | 52 (25.0) |
|  | 30-44 years |  | 52 (25.0) |
|  | 45-54 years |  | 52 (25.0) |
|  | 55-65 years |  | 52 (25.0) |
| Weight<sup>#</sup> | kg | 78 (66-87) |  |
| Hight<sup>#</sup> | m | 1.74 (1.68-1.80) |  |
| Body mass index | kg/cm<sup>2</sup> | 25.6 (22.8-28.1) |  |
| Body temperature | °C | 36 (35.7-36.2) |  |
| Systolic blood pressure | mmHg | 136 (121-145) |  |
| Diastolic blood pressure | mmHg | 78 (71-86) |  |
| Heart rate | beats/min | 78 (69-86) |  |
| Serum ApoA1 | mg/mL | 1.62 (1.49-1.79) |  |
| Serum ApoB | mg/mL | 0.90 (0.74-1.08) |  |
| Serum Insulin* | μU/mL | 13.2 (7.9-23.9) |  |
| Serum hsCRP | μg/mL | 0.74 (0.38-1.56) |  |
| Smoking<sup>#</sup> | No |  | 148 (71.2) |
|  | Yes |  | 60 (28.8) |
| Exercise intensity<sup>#</sup> | Moderate |  | 173 (83.2) |
|  | Intensive |  | 35 (16.8) |
| Exercise frequency<sup>#</sup> | <2-3x/week |  | 157 (75.5) |
|  | >2-3x/week |  | 51 (24.5) |
| Time passed since the last exercise<sup>#</sup> | <12 h |  | 13 (6.3) |
|  | 12-24 h |  | 23 (11.1) |
|  | 24-48 h |  | 14 (6.7) |
|  | >48 h |  | 158 (76.0) |

<sup>#</sup> self-reported data; \* in 34 samples values were below the limit of detection (3.06 μU/mL), only subjects with values larger than 3.06 μU/mL were included in the calculation of the median; ApoA1 – apolipoprotein A1; ApoB – apolipoprotein B; hsCRP – highly sensitive measurement of C-reactive protein.

Twenty-nine percent of study subjects were active smokers (30 males and 30 females). All study subjects reported regular exercise, with 16.8% describing it as intensive, however, for most (76%) the last exercise happened more than 48 hours before the blood draw. As expected, the median erythrocyte concentration in the blood of study subjects was the highest of all measured cell types at 4.91 x 10^9^/mL, with platelet and leukocyte median concentrations 22 times and 790 times lower, respectively. Neutrophils, the most numerous granulocytes, represented 56% of leukocytes in the blood, while lymphocytes and monocytes represented 33% and 8%, respectively (Supplementary Table 7). This distribution aligns with the typical composition of blood in apparently healthy adults (Troussard et al. 2014). To ensure consistency and minimise pre-analytical variables, plasma was prepared within 4 hours of blood draw, by the same individual at the same facility, following the standard protocol outlined in the Methods. Additionally, the high quality of plasma was confirmed by undetectable levels of erythrocytes and platelets on a routine haematology analyser before freezing. Furthermore, we analysed free haemoglobin concentrations in the thawed plasma samples, which were found to be well below the levels indicative of haemolysis, and assessed plasma PF4 levels as an indicator of platelet activation, which were 1468 (781-2370) ng/mL (Median (25-75%; Supplementary Table 8 and 9, Supplementary Figure 2; MIBlood-EV checklist).

To identify the cellular origins of circulating EVs in healthy adults, we next performed phenotyping of EV surface proteins by imaging flow cytometry directly in the defrosted blood plasma. Specifically, we determined concentrations of circulating EV subsets carrying one of the 25 studied markers of different blood or endothelial cells, delineated in Figure 2a and Supplementary Table 1. Some of the analysed markers also indicate precursor or stem cells. However, their percentage among circulating total nucleated cells in healthy adults at a steady state is less than 0.1% (Körbling and Anderlini 2001), so we inferred the precursor or stem cells are unlikely an important source of circulating EVs in our study. All investigated EV subsets, besides CD44+ EVs, were detected in the plasma of more than 95% of subjects, but showed a wide range of concentrations, from 4.2 (0.8-12) x 10^7^/mL to 1.9 (1.2-4.4) x 10^5^/mL (Median (25-75%); Figure 2b, Table 2). Circulating EV subsets with the highest median concentrations were positive for PAC-1, HLA-ABC, CD9, phosphatidylserine (PS) or CD227, while those with the lowest were positive for CD62P, CD90, CD13, CD14, CD44, or CD24.

**Figure 2:**
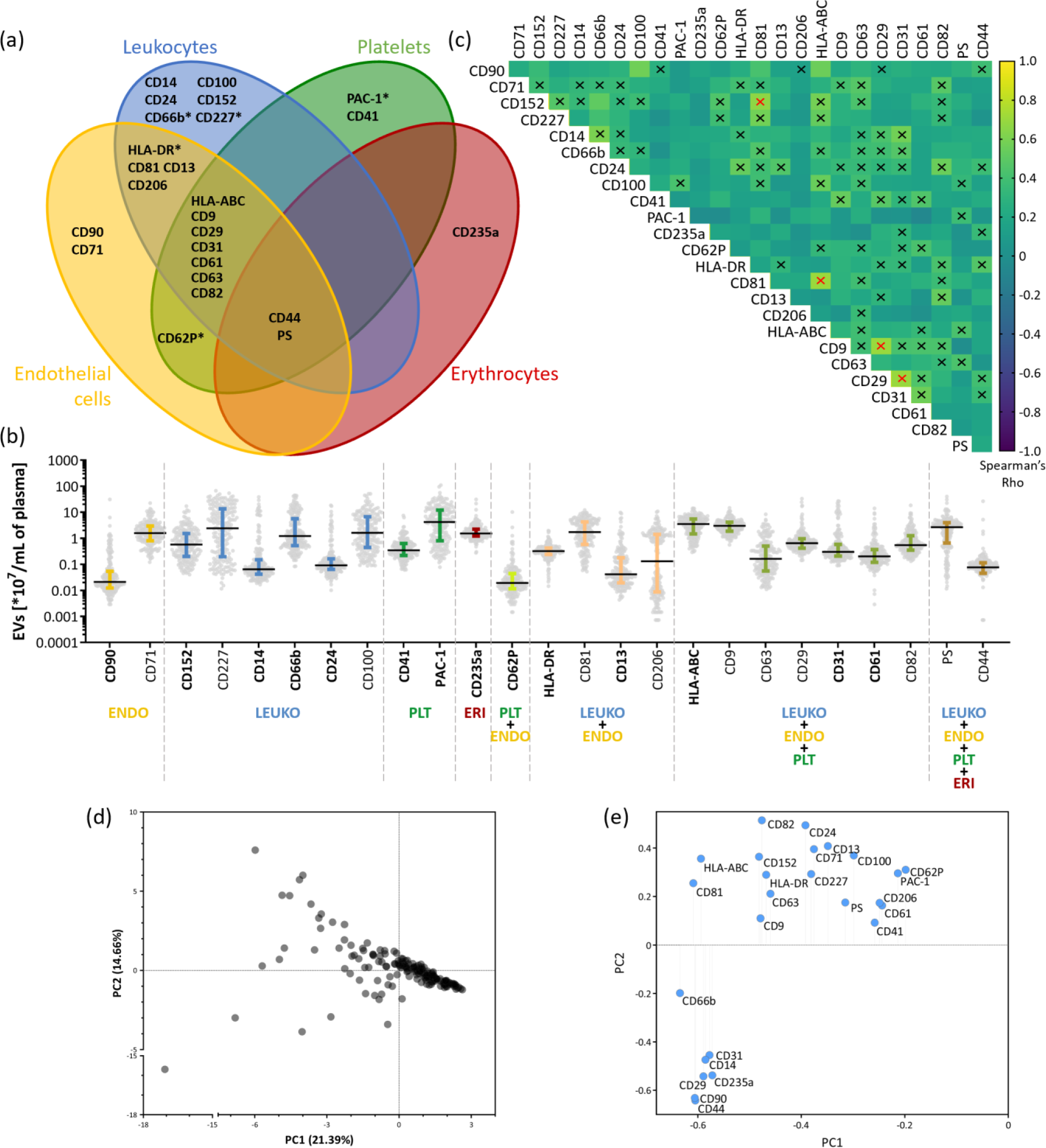
Quantification of circulating EV populations, with respect to their cell of origin, in the blood of healthy humans by imaging flow cytometry. **(a)** Representation of surface protein markers, included in imaging flow cytometry analysis, with respect to their presence on specific blood cell types. * indicates cell activation markers. **(b)** The measured concentrations of analysed circulating EV subsets, reported as median with interquartile range. Each dot represents one sample. Characteristic markers for the individual cell type are indicated in bold. ENDO – endothelial cells; LEUKO – leukocytes; PLT – platelets; ERI – erythrocytes. **(c)** Spearman’s correlations across all circulating EV subsets. Black cross – ρ: 0.3 – 0.59, Red cross – ρ ≥ 0.6. All depicted correlations are also statistically significant. **(d)** Principal component 1 vs 2 (PC1 vs. PC2) graph of the principal component analysis (PCA). Each dot represents one study subject. PC1 captures the most significant variation in the data, thus distances along the PC1 axis represent a larger variation than the same distances along the PC2 axis. **(e)** Loadings of original variables (circulating EV subset concentrations) of PCA. Circulating EV subsets with larger absolute values give the largest contribution to PC1 and PC2.

**Table 2:** Concentrations of EV subsets, carrying cell-specific surface markers, in the blood of healthy humans as measured by imaging flow cytometry.

| Marker | x 10<sup>6</sup>/mL of plasma<br>Median<br>(25-75%)<br>Total | x 10<sup>6</sup>/mL of plasma<br>Median<br>(25-75%)<br>Male | x 10<sup>6</sup>/mL of plasma<br>Median<br>(25-75%)<br>Female | % CV<br>Total<br>(male/female) | N<br>Total<br>(male/female) |
| --- | --- | --- | --- | --- | --- |
| CD90 | 0.21<br>(0.12-0.54) | 0.19<br>(0.12-0.42) | 0.26<br>(0.14-0.55) | 719 (215/663) | 200 (98/102) |
| CD71 | 15.80<br>(8.09-29.05) | 15.25<br>(7.85-27.48) | 16<br>(8.26-30.95) | 116 (102/123) | 200 (98/102) |
| CD152 | 5.77<br>(2.04-15.05) | 3.86<br>(1.63-14.05) | 7.86<br>(2.77-18.00) | 162 (131/163) | 200 (98/102) |
| CD227 | 24.10<br>(1.97-134.00) | 15.4<br>(1.52-130.50) | 32.1<br>(2.89-126.50) | 143 (146/141) | 200 (98/102) |
| CD14 | 0.64<br>(0.42-1.50) | 0.62<br>(0.39-0.97) | 0.76<br>(0.48-2.36) | 353 (334/349) | 200 (98/102) |
| CD66b | 12.25<br>(5.28-55.40) | 10.1<br>(4.38-40.68) | 13.45<br>(5.81-61.35) | 169 (175/163) | 200 (98/102) |
| CD24 | 0.91<br>(0.65-1.63) | 0.81<br>(0.57-1.32) | 1.05<br>(0.69-1.77) | 358 (413/319) | 200 (98/102) |
| CD100 | 16.25<br>(4.51-61.70) | 16.45<br>(3.28-57.73) | 16.25<br>(5.17-76.48) | 164 (161/161) | 200 (98/102) |
| CD41 | 3.44<br>(2.18-6.26) | 3.38<br>(2.01-6.10) | 3.59<br>(2.31-6.50) | 138 (137/138) | 200 (98/102) |
| PAC-1 | 41.85<br>(8.17-119.50) | 17.5<br>(5.75-75.75) | 62.9<br>(18.05-157.75) | 153 (153/139) | 200 (98/102) |
| CD235a | 15.40<br>(12.18-22.25) | 16.4<br>(12.23-25.50) | 15<br>(12.13-20.28) | 127 (102/150) | 200 (98/102) |
| CD62P | 0.19<br>(0.12-0.44) | 0.18<br>(0.12-0.43) | 0.22<br>(0.11-0.44) | 637 (201/560) | 200 (98/102) |
| HLA-DR | 3.22<br>(2.41-4.20) | 3.31<br>(2.34-3.82) | 3.13<br>(2.5-4.75) | 70 (64/74) | 200 (98/102) |
| CD81 | 17.30<br>(6.17-42.20) | 13.6<br>(5.07-39.08) | 26.15<br>(9.02-47.70) | 93 (104/83) | 200 (98/102) |
| CD13 | 0.42<br>(0.20-1.75) | 0.56<br>(0.24-1.98) | 0.34<br>(0.18-1.26) | 337 (359/295) | 200 (98/102) |
| CD206 | 2.15<br>(0.09-14.38) | 0.52<br>(0.07-13.48) | 7.19<br>(0.13-14.60) | 161 (183/122) | 197 (96/101) |
| HLA-ABC | 35.20<br>(14.68-54.45) | 27.95<br>(11.93-51.98) | 40.6<br>(17.28-65.60) | 74 (77/69) | 200 (98/102) |
| CD9 | 29.95<br>(18.70-41.25) | 27.45<br>(18.83-39.90) | 31.25<br>(17.65-42.90) | 63 (66/60) | 200 (98/102) |
| CD63 | 1.63<br>(0.57-5.04) | 1.03<br>(0.45-4.22) | 2.12<br>(0.78-5.52) | 182 (182/179) | 200 (98/102) |
| CD29 | 6.46<br>(4.17-9.50) | 6.41<br>(3.89-8.96) | 6.54<br>(4.87-11.18) | 254 (196/262) | 200 (98/102) |
| CD31 | 3.04<br>(2.11-5.81) | 2.91<br>(1.94-5.05) | 3.26<br>(2.29-6.89) | 260 (272/248) | 200 (98/102) |
| CD61 | 2.03 | 1.87 | 2.12 | 261 (241/281) | 199 (98/101) |

|  | (1.21-3.67) | (1.23-3.22) | (1.20-4.08) |  |  |
| --- | --- | --- | --- | --- | --- |
| **CD82** | 5.46<br>(3.52-12.35) | 5.94<br>(3.68-14.95) | 5.3<br>(3.27-9.33) | 159 (160/157) | 200 (98/102) |
| **PS** | 26.75<br>(6.79-39.50) | 20.85<br>(4.61-36.13) | 29.95<br>(11.55-41.70) | 136 (173/99) | 200 (98/102) |
| **CD44** | 0.90<br>(0.64-1.23) | 0.96<br>(0.70-1.23) | 0.8<br>(0.62-1.28) | 680 (132/613) | 169 (77/92) |

Markers, shaded in grey, are the most characteristic markers of the individual cell type.

To estimate the relative abundance of circulating EV subsets with respect to their cell origin, we used established markers for the immunophenotyping by flow cytometry (Kalina et al. 2019; Grant et al. 2021; de Oliveira et al. 2023; Berckmans et al. 2019; Spurgeon and Frelinger 2022). We designated CD235a+ subsets as erythrocyte EVs, CD41+ and PAC1+ subsets as platelet EVs, and CD61+, CD31+, and CD62P+ subsets as originating from platelets or endothelial cells. Additionally, CD90+ EVs were identified as originating from activated endothelial cells in our study (Wandel et al. 2012), as we do not expect precursor or stem cells to be an important source of circulating EVs (Körbling and Anderlini 2001). Among leukocytes, CD66+ EVs originate from activated granulocytes, CD152+ EVs from T lymphocytes, and CD24+ EVs from B lymphocytes or granulocytes. Antigen-presenting cells (APCs) were associated with HLA-DR+ EVs, while CD14+ subsets were identified as monocyte EVs.

When considering the median concentrations of cell-specific EV subsets (Figure 2b, Table 2), activated platelet EVs emerged as the most abundant among circulating EVs, followed by erythrocyte EVs, granulocyte EVs, T lymphocyte EVs, EVs from APCs (including monocyte EVs), and endothelial EVs in the lowest concentration. We investigated the relationship between circulating EV subset concentrations and their corresponding cells-of-origin, as determined by complete blood count (Supplementary Table 7). No statistically significant correlation was found. Notably, both the highest (PAC-1+) and lowest (CD62P+) median concentrations of circulating EV subsets originated from activated platelets. To further explore possible patterns among EV subsets, we assessed correlations among all 25 EV subset concentrations (Figure 2c). We identified only four moderate associations (ρ > 0.6, marked with a red cross in Figure 2c), suggesting diverse patterns of EV subset release and/or uptake, even from the same cell type.

Furthermore, we performed the principal component analysis (PCA) to independently identify similarities among study subjects based on their circulating EV subsets concentrations. Principal components PC1 and PC2 revealed a compact cluster of study subjects, suggesting high multidimensional similarity in their circulating EV subset concentrations. The remaining study subjects appeared scattered, lacking distinct patterns or clusters (Figure 2d). Loadings in PCA represent the contribution of each variable to each principal component, revealing the most influential variables in explaining data variance. In our analysis, granulocyte-derived CD66b+ EVs, CD81+ EVs, activated endothelial CD90+ EVs, and CD44+ EVs contributed most to the variability observed in the PC1 component (Figure 2e). EV subsets derived mostly from platelets (CD41+, CD61+, PAC-1+, and CD62P+ EVs) formed a distinct cluster in the loadings graph of PCA, yet their contribution to PC1 variability was minimal. Additionally, among three of the most commonly studied EV-related tetraspanins, CD9+ and CD63+ EVs showed similar loadings on PC1, distinct from the CD81+ EVs.

Altogether, we described the physiological EV population composition with respect to their cellular source in the blood of healthy humans. EV subset concentrations did not directly reflect concentrations of blood cells, indicating diverse patterns of EV subset release and/or uptake, even for EVs originating from the same cell type.

### Tetraspanin+ circulating EVs largely originate from platelets and lymphocytes in the blood of healthy humans

We detected high median concentrations of circulating EV subsets, positive for tetraspanins CD9 and CD81, while CD63+ EV concentration was much lower. To investigate the cellular origin of those tetraspanin+ circulating EVs, we first enriched EVs from the blood plasma of all study subjects using established sucrose cushion ultracentrifugation (Figure 1; (Holcar et al. 2020)). As evaluated by NTA, the median (25-75%) mean concentration of particles in enriched circulating EVs across study subjects was 5.65 x 10^9^ (5.07–6.19 x 10^9^) particles/mL of plasma, with median (25-75%) modal diameter of 151.7 (144.8-158.9) nm (Supplementary Figure 3a, b). At the same time, ELISA showed a substantial decrease of ApoA1 and ApoB levels after enrichment of EVs from the subject’s plasma (Supplementary Figure 2; 0.0057% and 0.0017% (with 92 samples below the limit of detection) of ApoA1 and ApoB, respectively). A statistically significant difference in the NTA detected particle concentrations was observed between sexes (5.74 x 10^9^ and 5.44 x 10^9^ particles/mL for males and females, respectively; p = 0.01715, Supplementary Figure 3c). However, this difference was within the calculated analytical CV for NTA concentration measurements of 7.89%, equivalent to 0.45 x 10^9^ particles/mL (Supplementary Figure 3d). Therefore, it might be attributed to the imprecision inherent in NTA concentration measurements rather than holding biological importance.

We next performed phenotyping of EVs based on their expression of 37 surface proteins with semi-quantitative multiplex bead-based flow cytometry on enriched circulating EVs across study subjects (Figure 1). Specifically, we determined normalised median fluorescence intensity (MFI) as a measure of the expression level of cell-type specific protein markers (delineated in Figure 3a, Supplementary Table 10), present on the surface of circulating CD9 and/or CD81 and/or CD63 positive (tetraspanin+) EVs (Figure 3). Twenty-three markers were detected in the plasma of more than 91% of subjects and included in further analysis, showing up to a 17-fold difference in the level of expression on the tetraspanin+ circulating EVs (from 19.63 (11.59-29.21) to 1.13 (0.85-1.66) normalised median MFI (25-75%); Figure 3b, Table 3). Thirteen surface protein markers were excluded from the analysis due to low detection across study subjects (measured MFIs were in the range of or under the limit of detection), while ROR1 surface protein was excluded as it is a cancer marker.

**Figure 3:**
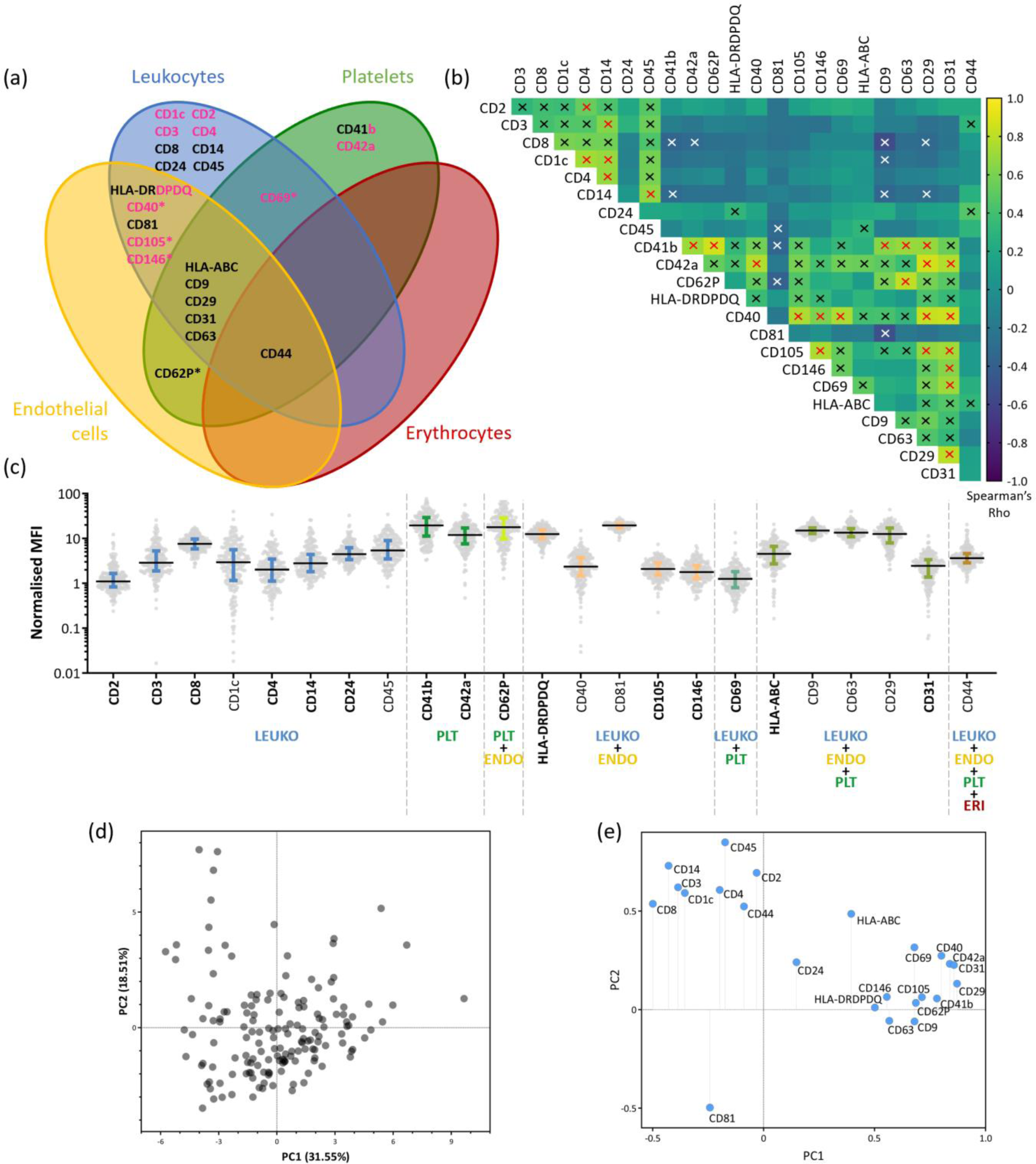
Tetraspanin+ circulating EVs largely originate from platelets and lymphocytes in the blood of healthy humans, as determined by multiplex bead-based flow cytometric analysis. **(a)** Representation of surface protein markers, included in multiplex bead-based flow cytometric analyses, with respect to their presence on specific blood cell types; * indicates cell activation markers. Black text indicates markers were also included in the imaging flow cytometry analysis, while those in pink were only analysed by multiplex bead-based FC. **(b)** Normalised median fluorescence intensity (MFI) of tetraspanin+ circulating EVs expressing one of the studied markers, reported as median with interquartile range. Each dot represents one sample. Characteristic markers for the individual cell type are indicated in bold. ENDO – endothelial cells; LEUKO – leukocytes; PLT – platelets; ERI – erythrocytes. **(c)** Spearman’s correlation across all tetraspanin+ circulating EVs expressing specific markers, Black cross – ρ: 0.3 – 0.59, Red cross – ρ ≥ 0.6. White cross – ρ ≤ -0.3. All correlations were statistically significant. **(d)** Principal component 1 vs 2 (PC1 vs. PC2) graph of the principal component analysis (PCA). Each dot represents one study subject. PC1 captures the most significant variation in the data, thus distances along the PC1 axis represent a larger variation than the same distances along the PC2 axis. **(e)** Loadings of original variables (circulating EV subset concentrations) of PCA. Tetraspanin+ circulating EV subsets with larger absolute values give the largest contribution to PC1 and PC2.

**Table 3:** Expression levels of cell-specific markers on tetraspanin+ circulating EV in the blood of healthy humans as measured by multiplex bead-based flow cytometry.

| Marker | Normalised MFI<br>Median (25-75%)<br>total | Normalised MFI<br>Median (25-75%)<br>male | Normalised MFI<br>Median (25-75%)<br>female | % CV<br>Total<br>(male/female) | N<br>Total<br>(male/female) |
| --- | --- | --- | --- | --- | --- |
| CD2 | 1.13<br>(0.85-1.66) | 1.09<br>(0.78-1.61) | 1.21<br>(0.9-1.94) | 91 (63/100) | 191 (95/96) |
| CD3 | 2.91<br>(1.93-5.33) | 3.08<br>(1.95-5.37) | 2.85<br>(1.85-5.33) | 87 (90/83) | 199 (99/100) |
| CD8 | 7.61<br>(5.93-9.75) | 7.81<br>(6.25-9.47) | 7.13<br>(5.79-9.8) | 38 (37/38) | 205 (103/102) |
| CD1c | 3.20<br>(1.45-6.15) | 3.2<br>(1.68-5.95) | 2.98<br>(1.18-6.24) | 107 (90/119) | 189 (92/97) |
| CD4 | 2.05<br>(1.18-3.51) | 2.06<br>(1.21-3.13) | 2.03<br>(1.15-3.78) | 111 (79/127) | 196 (97/99) |
| CD14 | 2.85<br>(1.85-4.41) | 2.97<br>(2.04-4.56) | 2.66<br>(1.65-4.35) | 93 (90/97) | 202 (100/102) |
| CD24 | 4.46<br>(3.39-6.10) | 4.25<br>(3.38-5.75) | 4.69<br>(3.44-6.84) | 64 (56/69) | 205 (102/103) |
| CD45 | 5.48<br>(3.52-90) | 5.3<br>(3.49-9.62) | 5.73<br>(3.61-8.33) | 67 (63/71) | 205 (102/103) |
| CD41b | 19.63<br>(11.59-29.21) | 20.06<br>(11.16-28.58) | 19.39<br>(11.77-29.52) | 61 (62/60) | 205 (103/102) |
| CD42a | 12.00<br>(7.86-17.22) | 11.95<br>(7.54-16.43) | 12.85<br>(8.24-17.84) | 57 (62/53) | 205 (103/102) |
| CD62P | 17.9<br>(10.08-28.93) | 17.55<br>(8.22-27.2) | 18.35<br>(12.75-30.52) | 72 (79/66) | 205 (103/102) |
| HLA-DRDPDQ | 12.6<br>(10-15.48) | 12.6<br>(10.63-15.65) | 12.55<br>(9.86-15.21) | 32 (32/32) | 205 (103/102) |
| CD40 | 2.42<br>(1.59-3.85) | 2.45<br>(1.55-3.69) | 2.39<br>(1.62-4.06) | 67 (74/59) | 196 (100/96) |
| CD81 | 19.5<br>(17.32-21.54) | 19.5<br>(17.52-21.18) | 19.5<br>(17.3-21.79) | 16 (17/16) | 206 (103/103) |
| CD105 | 2.13<br>(1.54-2.87) | 2.28<br>(1.62-3.07) | 2.07<br>(1.44-2.63) | 43 (41/45) | 197 (100/97) |
| CD146 | 1.79<br>(1.29-2.48) | 1.75<br>(1.26-2.36) | 1.9<br>(1.3-2.53) | 51 (52/50) | 196 (99/97) |
| CD69 | 1.26<br>(0.8-1.81) | 1.26<br>(0.79-1.81) | 1.26<br>(0.8-1.79) | 85 (97/69) | 195 (99/96) |
| HLA-ABC | 4.63<br>(3.09-6.87) | 4.65<br>(3.32-6.78) | 4.62<br>(2.75-6.91) | 57 (63/49) | 194 (99/95) |
| CD9 | 15.08<br>(12.94-17.34) | 15.05<br>(12.79-17.11) | 15.1<br>(13.2-17.73) | 24 (26/22) | 206 (103/103) |
| CD63 | 13.59<br>(11.05-16.55) | 12.4<br>(10.57-15.69) | 14.36<br>(11.8-17.42) | 36 (39/33) | 206 (103/103) |
| CD29 | 12.63<br>(7.98-17.2) | 12.66<br>(7.74-17.05) | 12.63<br>(8.95-17.28) | 49 (52/47) | 206 (103/103) |
| CD31 | 2.51 | 2.41 | 2.59 | 60 (67/53) | 196 (99/97) |

|  | (1.51-3.38) | (1.37-3.33) | (1.73-3.39) |  |  |
| --- | --- | --- | --- | --- | --- |
| **CD44** | 3.65<br>(2.88-4.61) | 3.68<br>(3.03-4.82) | 3.48<br>(2.82-4.57) | 59 (64/52) | 205 (102/103) |

#Values above the control limit. Markers, shaded in grey are the most characteristic markers of representative cell type.

Tetraspanin+ circulating EVs showed the highest expression levels for platelet markers CD41b, CD62P (also a marker of activated endothelium) and CD42a, alongside the more general marker CD29 and leukocyte protein HLA-DRDPDQ. Conversely, expression was the lowest for lymphocyte markers CD2 and CD69, and the endothelial marker CD146. Other evaluated markers with intermediate expression were leukocyte markers CD8 (lymphocyte T), CD45, CD24 (lymphocyte B or granulocytes), CD3 (lymphocyte T), CD14 (monocytes), CD105 (macrophages and endothelium) and CD4 (mostly lymphocyte T), ranked from highest to lowest. Additionally, more general markers of blood cells, such as HLA-ABC (present on all nucleated blood cells), CD44, CD1c, CD31 and CD40, demonstrated intermediate expression on tetraspanin+ circulating EVs (Figure 3b, Table 3). Erythrocyte-specific markers are not included in this multiplex bead-based flow cytometry analysis. Additional analyses of correlations between different markers of activated platelets (CD62P+ EVs, PF4 concentration) and concentration of platelets in whole blood or plasma excluded non-physiological platelet activation after the blood draw (Supplementary Figure 2). Importantly, examination of normalised expression levels (measured by multiplex bead-based flow cytometry) or normalised concentrations (measured by imaging flow cytometry) of markers CD14, CD24, CD41, CD62P, HLA-DR, HLA-ABC, CD29, CD31, and CD44 on circulating EVs revealed distinct profiles for tetraspanin+ circulating EVs compared to the overall circulating EV population (Supplementary Figure 4). Thus, tetraspanin+ circulating EVs are not representative of the entire circulating EV population in healthy human blood.

To determine if markers specific to a particular cell type exhibited similar expression patterns on tetraspanin+ circulating EVs, we evaluated correlations among the 23 studied surface proteins (Figure 3c) and found 26 pairs with moderate correlations (ρ > 0.6, red cross). Among those CD41b expression correlated with CD42a, CD62P, CD9 and CD63 (ρ > 0.6, red cross), indicating a conserved pattern of marker expression on CD9+ and/or CD63+ EVs released from platelets across study subjects. Similarly, the expression of CD105 correlated with CD146, CD31 and CD29, indicating a conserved pattern of marker expression on tetraspanin+ circulating EVs released from endothelium. For leukocyte markers, we mostly observed individual correlations (e.g. CD2 with CD4; CD14 with CD3, CD4 and CD45), which might reflect the diversity of leukocyte cell types.

We conducted PCA to assess similarities among study subjects based on the expression levels of all studied surface markers on tetraspanin+ circulating EVs (Figure 3d). Subjects were less tightly clustered when compared to the PCA analysis of circulating EV subset concentrations (Figure 2d). Expression levels of markers CD29 and CD31 (both cell-type nonspecific), followed by CD42a and CD40 (typical for B lymphocytes, macrophages, and dendritic cells) on tetraspanin+ circulating EVs, influenced most of the variability observed in the PC1 component (Figure 3e). CD9+ and CD63+ EVs showed similar loadings on PC1, while CD81+ EVs diverged significantly, indicating distinct cell origins for the latter.

In summary, tetraspanin+ circulating EVs in healthy humans largely originate from platelets and to a lesser extent from lymphocytes, while some also originate from other leukocyte types or endothelium. Interestingly, platelet and endothelial markers show a conserved pattern of expression on tetraspanin+ circulating EVs across study subjects.

### Circulating EVs show high inter-individual biological variability in a healthy human cohort

While high inter-individual biological variation is commonly observed in established plasma analytes, it has yet to be studied for circulating EVs. To assess inter-individual biological variation in the concentration of circulating EV subsets, positive for one of the 25 studied markers of different blood cells or endothelial cells, we calculated the CV across all study subjects (Figure 4a, Table 2). Inter-individual variation was high for all circulating EV subset concentrations (median CV across all EV subsets was 162%) but varied a lot between different EV subsets. Circulating CD90+, CD44+ and CD62P+ EV concentrations had a CV of 719%, 680% and 637%, respectively, while CD9+, HLA-DR+, HLA-ABC+ and CD81 EV concentrations were more consistent across study subjects (63%, 70%, 74% and 93%, respectively). Interestingly, high inter-individual variability observed for circulating CD90+, CD44+ and CD62P+ EVs was mostly at the expense of high CVs observed in females (663%, 613% and 560%, respectively, Supple. Figure 5a). CV was inconsistent across different EV subsets originating from the same cell type, and independent of activation state across various cell types. For example, CD41+, PAC-1+ and CD62P+ EV subsets originating from platelets had CVs of 138%, 153% and 637%, respectively. Similarly, CD90+, CD66b+, PAC-1+ and CD62P+ EV subsets, originating from activated endothelial cells, granulocytes and platelets, respectively, had CVs in the range of 719% to 153%.

**Figure 4:**
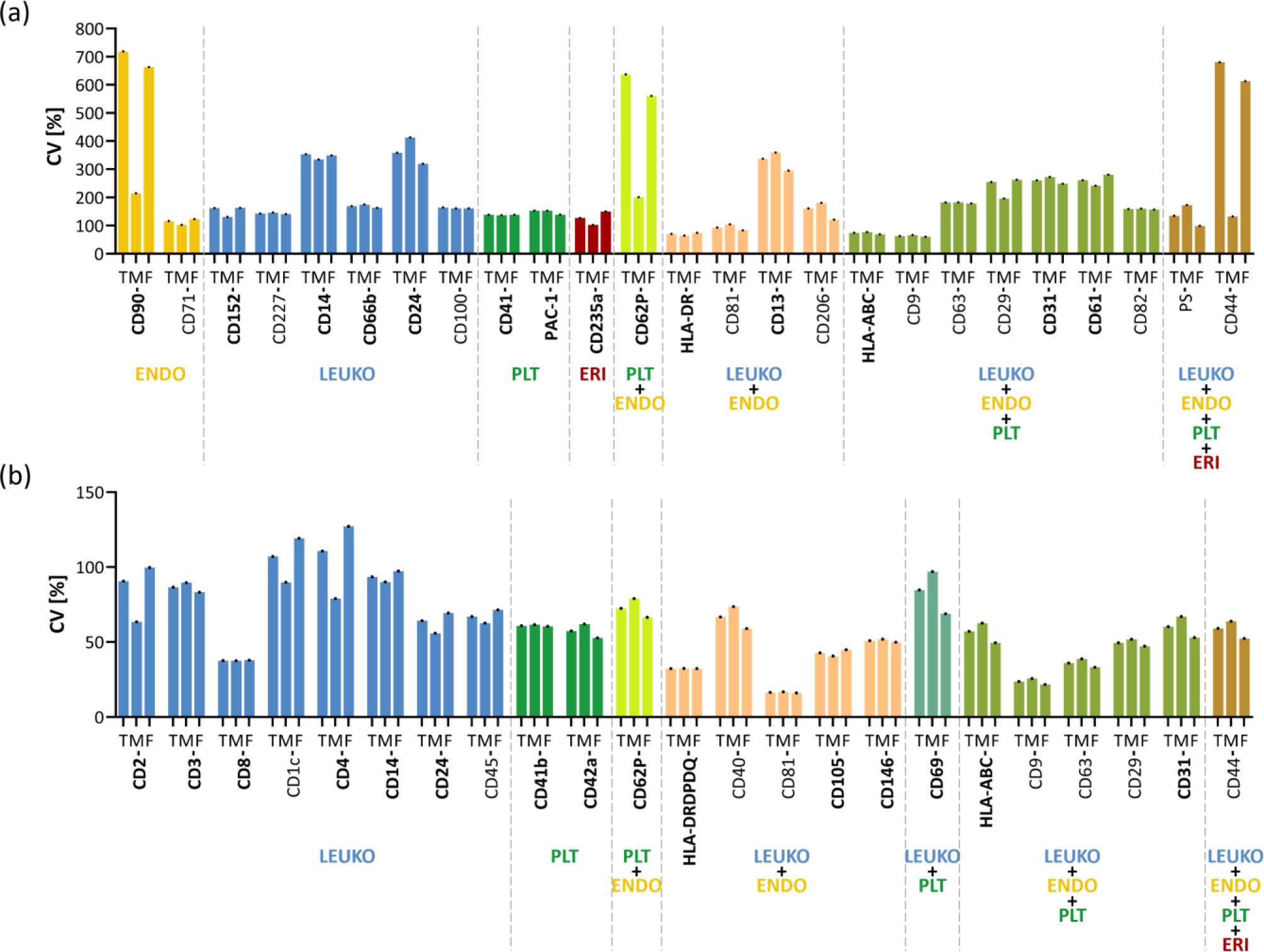
Analysis of inter-individual biological variability of circulating EV subset concentrations and tetraspanin+ circulating EV marker expression levels. Coefficient of variations of **(a)** circulating EV subset concentrations and **(b)** normalised MFI for markers on tetraspanin+ circulating EVs, across study subjects. CV – coefficient of variation; T-total; M – male; F – Female; ENDO – endothelial cells; LEUKO – Leukocytes; PLT – platelets; ERI – erythrocytes.

**Figure 5:**
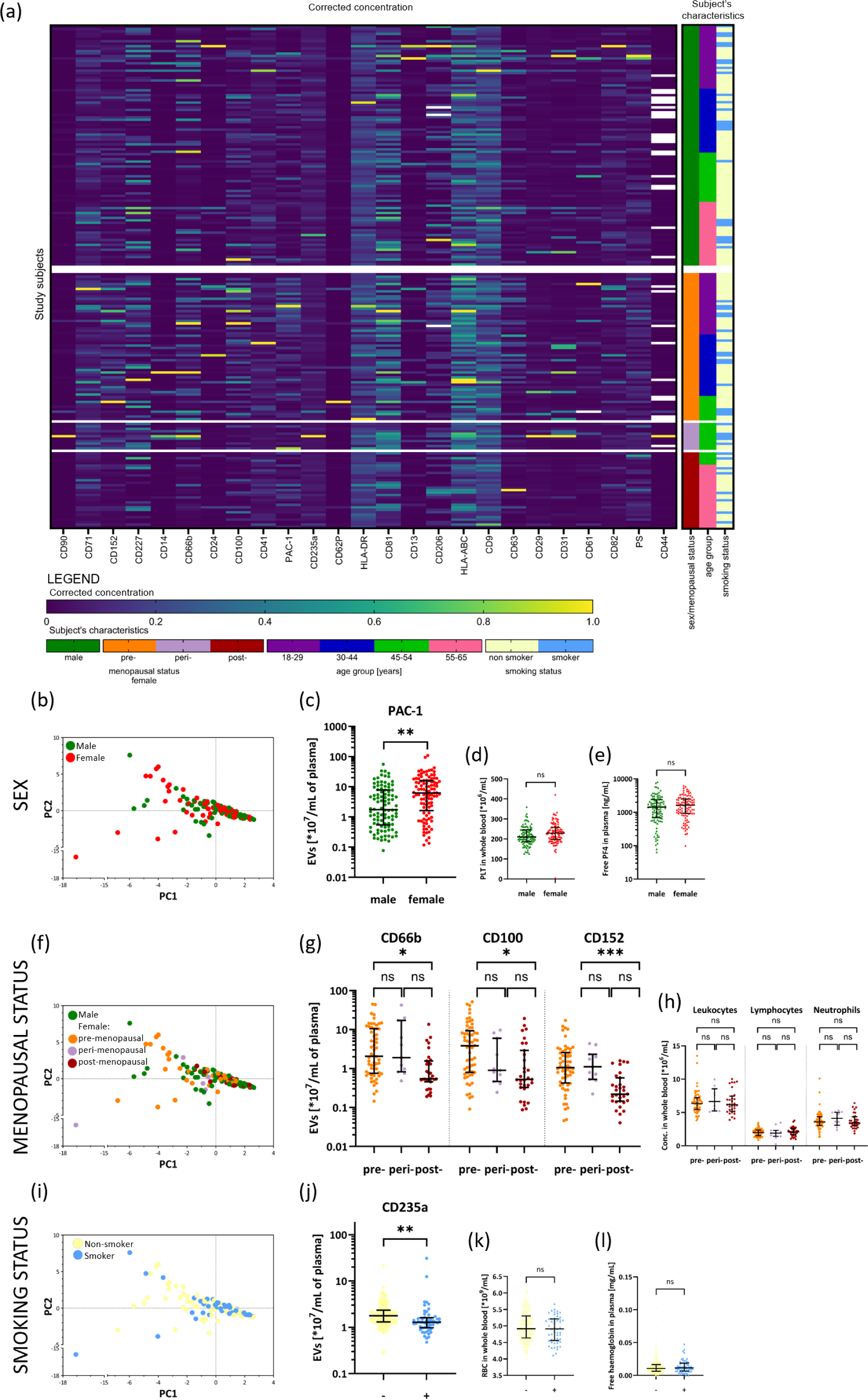
Sources of inter-individual biological variability observed in circulating EV subsets. **(a)** Results of imaging flow cytometry analysis of circulating EV subsets, normalised to the largest measured value of each circulating EV subset. Each line represents one study subject. Subjects are arranged by their increasing age, sex, menopausal status and smoking status. **(b)** PCA plot, coloured according to the sex of study subjects. Influence of sex on **(c)** concentration of PAC-1+ circulating EVs in plasma, **(d)** concentration of platelets in whole blood and **(e)** concentration of free PF4 in plasma. **(f)** PCA plot, coloured according to the menopausal status of study subjects. Influence of menopause status on concentrations of **(g)** CD66b+, CD100+ and CD152+ circulating EVs in plasma, and **(h)** concentration of leukocytes, lymphocytes and neutrophils in whole blood. **(i)** PCA plot, coloured according to the smoking status of study subjects. Influence of smoking on **(j)** concentration of CD235a+ circulating EVs in plasma, **(k)** concentration of erythrocytes in whole blood, and **(l)** concentration of free haemoglobin in plasma. Each dot represents one study subject. Black lines represent the median with the interquartile range. * - p < 0.002; ** - p < 0.0004; *** - p < 0.00004; ns – nonsignificant; PLT – platelets, PF4 – plasma factor 4; RBC – erythrocytes; WBC - leukocytes.

The expression level (normalised median MFI) of 23 cell-type specific protein markers on the surface of tetraspanin+ circulating EVs, was much more consistent across study subjects (median CV across all EV subsets was 60%; Figure 4b, Table 3) compared to the concentration of circulating EV subsets. Still, higher inter-individual variability was observed for CD4, CD1c, CD14 and CD2 expression on tetraspanin+ circulating EVs (CVs of 111%, 107%, 93% and 91%, respectively), while the lowest was observed for HLA-DRDPDQ, CD8, and CD105 (CVs of 32%, 38% and 43%, respectively). Again, high inter-individual variability was driven by higher CVs in females (127% for CD4, 119% for CD1c and 100% for CD2, Figure 4b, Table 3 and Supplementary Figure 5b).

Taken together, circulating EV subset concentrations show high inter-individual biological variability in healthy humans, with differing levels of variability for EV subsets originating from the same cell type or with respect to its activation state. In comparison, the tetraspanin+ subpopulation of circulating EVs shows lower variation in expression levels of cell-type specific protein markers.

### Sex, menopausal status and smoking contribute to inter-individual biological variability of specific circulating EV subsets, originating from platelets, leukocytes or erythrocytes

Our study of inter-individual variability of circulating EV subset concentrations indicated sex could be one of the underlying factors of this variability. To comprehensively investigate the sources of observed biological variability, we collected data on the following demographic (age, sex), clinical (height, weight, body mass index, blood pressure, blood pulse, menopause status, complete blood count and lipoprotein concentrations), and lifestyle (exercise regimen, time since last exercise, smoking status) factors across the study subjects (Table 1) and analysed their association with measured circulating EV subset concentrations.

The heatmap visually represents all measured circulating EV subset concentrations for every study subject. We arranged them by increasing age, sex, menopausal status, and smoking (Figure 5a). There is a visible pattern of higher EV concentrations for most EV subsets in pre- and peri-menopausal females, compared to post-menopausal females or males. Importantly, the PCA showed none of the analysed subject’s factors significantly influenced the measured concentrations of the 25 circulating EV subsets on a global level. However, we identified sex, menopausal status, and smoking as factors associated with specific circulating EV subset concentrations (Figure 5b-l).

Although sex did not result in distinct groupings in PCA of EV subset concentrations (Figure 5b), platelet PAC-1+ EVs, exhibited significantly higher concentrations in females compared to males (p = 0.00010, Figure 5c). This could not be attributed to differences in platelet concentrations in whole blood or free PF4 levels in plasma, as no significant differences were observed between females and males (Figure 5d, e). When data represented on the PCA graph was colour-coded by menopausal status, no distinct patterns were observed (Figure 5f). However, CD66b+, CD100+, and CD152+ EVs, originating from granulocytes, leukocytes, and T lymphocytes, respectively, exhibited significantly higher concentrations in pre-menopausal females compared to post-menopausal ones (p = 0.00109, p = 0.00001, and p = 0.00068, respectively; Figure 5g). Notably, blood leukocyte, lymphocyte, or neutrophil concentrations were not dependent on female menopausal status (Figure 5h). Finally, although smoking did not influence the overall concentration of circulating EV subsets according to PCA (Figure 5i), CD235a+ EVs originating from erythrocytes showed significantly lower concentrations in smokers compared to non-smokers (p = 0.00009, Figure 5j). This could not be explained by the influence of smoking on blood erythrocyte levels or haemolysis, since no significant differences were observed among our study subjects (Figure 5k, l).

To conclude, no global influence on studied circulating EV subsets was observed for any of the subject’s factors. However, sex, menopausal status and smoking were shown to contribute to inter-individual biological variability of specific circulating EV subsets, originating from platelets, leukocytes or erythrocytes, respectively.

### Smoking and age contribute to inter-individual biological variability of specific tetraspanin+ circulating EVs, originating from platelets, leukocytes or endothelial cells

The tetraspanin+ subpopulation of circulating EVs exhibited lower biological variation in the expression levels of cell-type specific protein markers, in comparison to the observed variation in circulating EV subset concentrations. However, their CVs still ranged from 32% to 111%, prompting us to investigate the factors contributing to this variability. The heatmap visually represents all measured expression levels of markers on tetraspanin+ circulating EVs for every study subject, which were arranged by increasing age, sex, menopausal status, and smoking (Figure 6a). We next examined the contribution of individual subject factors (Table 1) to variability in the expression levels of cell-type specific protein markers on tetraspanin+ circulating EVs. Again, none of the analysed subjects’ factors significantly influenced the observed variability on a global level. However, association analysis revealed that smoking and age were correlated with the expression levels of specific protein markers present on the surface of tetraspanin+ circulating EVs (Figure 6b-j).

**Figure 6:**
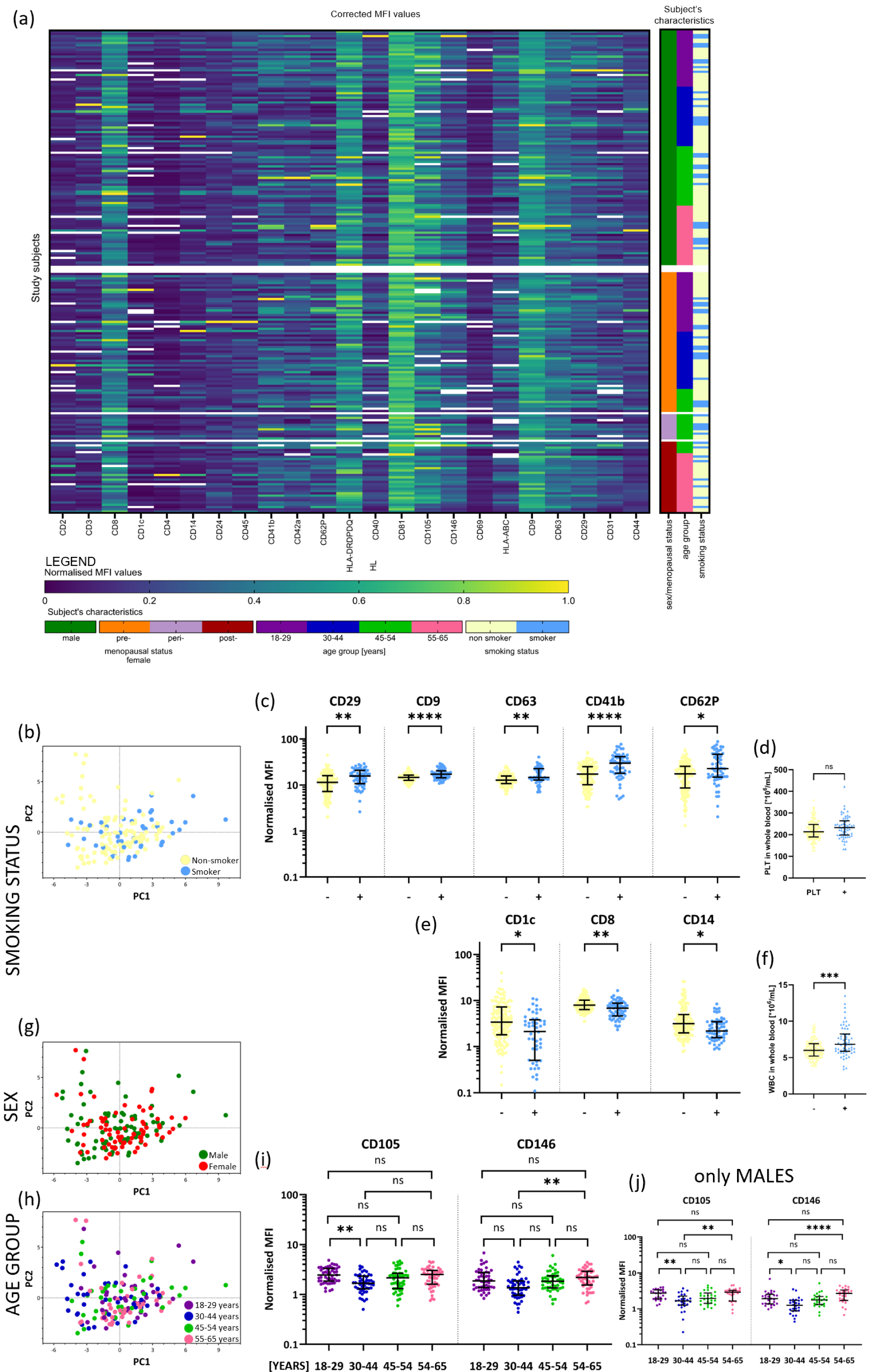
Sources of inter-individual biological variability observed in tetraspanin+ circulating EVs expressing specific protein markers. **(a)** Results of multiplex bead-based flow cytometry analysis of tetraspanin+ circulating EVs, corrected by dividing normalised MFIs by the largest measured normalised MFI value of each marker on tetraspanin+ circulating EVs. Each line represents one study subject. Subjects are arranged by their increasing age, sex, menopausal status and smoking status. **(b)** PCA plot, coloured according to the smoking status of study subjects. Influence of smoking status on **(c)** expression levels of CD29+, CD9+, CD63+, CD41b+ and CD62P+ tetraspanin+ circulating EVs in plasma, and **(d)** platelet concentration in whole blood. PLT – platelets. Influence of smoking status on **(e)** expression levels of CD1c+, CD8+ and CD14+ tetraspanin+ circulating EVs in plasma, and **(f)** leukocyte concentration in whole blood. WBC – leukocytes. PCA plots, coloured according to **(b)** sex and **(h)** age group of study subjects. **(i)** Influence of age on expression levels of CD105+ and CD164+ tetraspanin+ circulating EVs. **(j)** Influence of age on expression levels of CD105+ and CD164+ tetraspanin+ circulating EVs presented only in man. Each dot represents one study subject. Black lines represent median with interquartile range; * - p < 0.002; ** - p < 0.0004; *** - p < 0.00004; **** - p < 0.000004; ns – nonsignificant; MFI – median fluorescence intensity.

While smoking did not show distinct clusters in PCA (Figure 6b), it did affect the expression levels of eight protein markers on tetraspanin+ circulating EVs. Median normalised MFI levels of five of these markers (CD29, a nonspecific cell marker; CD41b and CD62P, predominantly originating from platelets; and CD9 and CD63, which also appeared to originate primarily from platelets) were significantly increased in smokers (p = 0.000162, p = 0.000001, p = 0.000821, p = 0.000002, and p = 0.000248, respectively; Figure 6c), despite no statistically significant difference in blood platelet concentration between smokers and non-smokers (Figure 6d). Conversely, median normalised MFI levels of CD1c, as well as T lymphocyte-derived CD8 and monocyte-derived CD14 markers on tetraspanin+ circulating EVs, were decreased in smokers (p = 0.000535, p = 0.000227, and p = 0.00797, respectively; Figure 6e), despite a significant increase in their blood leukocyte concentration (p = 0.0001, Figure 6f). While neither sex nor age group revealed any distinct patterns in PCA (Figure 6g, h), age affected the expression levels of CD146 and CD105 present on tetraspanin+ circulating EVs originating from endothelial cells (CD105 also from T lymphocytes; Figure 6i). Median normalised MFI levels of CD105 and CD146 markers on tetraspanin+ circulating EVs were significantly higher in the 18 – 29 years age group and 54 – 65 years age group, respectively, compared to the 30 – 44 years age group (p = 0.000306 and p = 0.000332, respectively). The difference was even more pronounced when analysing only males, indirectly suggesting the influence of sex on expression levels of CD146 and CD105 on tetraspanin+ circulating EVs (Figure 6j).

Altogether, while no global influence on studied circulating EV subsets was observed for any of the subject’s factors, smoking and age were shown to contribute to inter-individual biological variability of tetraspanin+ subpopulation of circulating EVs originating from platelets, leukocytes or endothelial cells.

## Discussion

This study comprehensively quantified EVs with respect to cell origin in the blood of a large cohort of healthy humans, to improve understanding of the physiological circulating EV population, its biological variability and sources of observed variability (Figure 8). As revealed by imaging and multiplex bead-based flow cytometry analyses, cell-specific circulating EV subsets show a wide range of concentrations, which do not directly reflect concentrations of blood cells, indicating diverse patterns of EV subset release and/or uptake, even for EVs originating from the same cell type. Interestingly, the tetraspanin+ subpopulation of circulating EVs largely originates from platelets and to a lesser extent from lymphocytes. We provide the first proof of high inter-individual biological variability of circulating EV subset concentrations in healthy humans, with differing levels of variability for EV subsets originating from the same cell type or with respect to its activation state. Tetraspanin+ circulating EVs show lower variation in expression levels of cell-type specific protein markers. Sex, menopausal status, age and smoking contribute to inter-individual biological variability observed in concentrations of specific circulating EV subsets or specific protein marker expression levels on tetraspanin+ EVs, respectively. Nevertheless, no significant global effect of the subjects’ factors on the circulating EVs or its tetraspanin+ subpopulation was identified.

**Figure 8:**
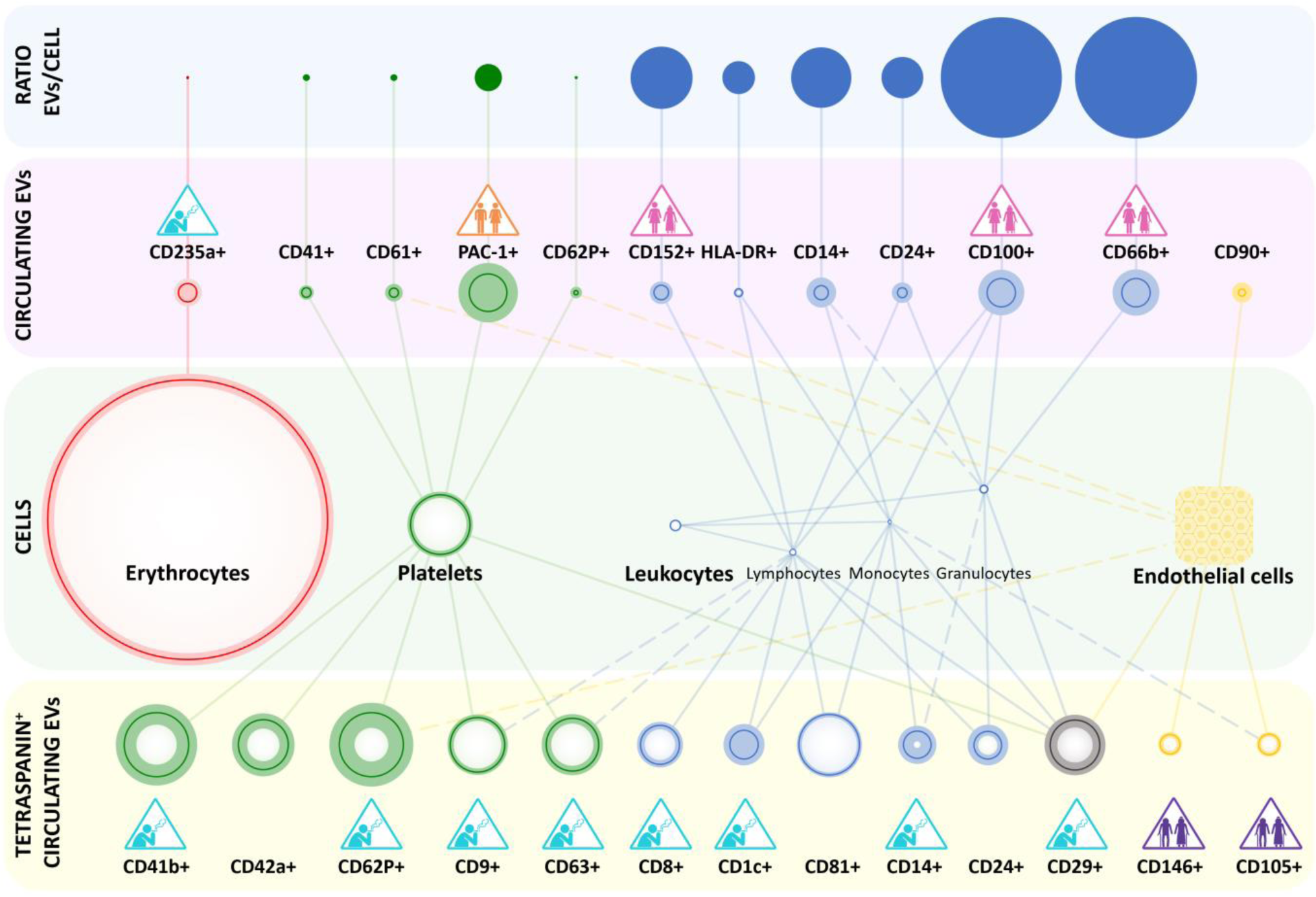
Schematic representation of circulating EVs in healthy humans: cellular origin and biological variability. Cell-specific circulating EV subsets show a wide range of concentrations as measured by imaging flow cytometry (pink field), which do not directly correspond to the concentrations of blood cells (green field). The sizes of the circles directly reflect the measured relative average concentrations of blood cells per mL of blood or EVs per mL of plasma, while the biological variability determined by the coefficient of variation (CV) is indicated by the shading of the circle borders. For endothelial cells, the size of the symbol does not reflect concentration as their count is unknown. Cellular sources of specific circulating EV subsets are indicated by a solid line from cells to EVs. When multiple cell types contribute to EVs, the most frequent cellular source is depicted with a solid line, while less common or less probable sources are shown with dashed lines. The estimated number of circulating EVs, carrying the indicated protein marker, per the parental blood cell type (blue field) suggests diverse patterns of EV subset release and/or uptake, even for EVs originating from the same cell type. The sizes of the full circles represent the relative ratio of EVs/cell. The EV-to-cell ratio for all EVs originating from any leukocytes is calculated as the number of EVs per leukocyte. The tetraspanin+ subpopulation of circulating EVs shows lower biological variation in expression levels of cell-type specific protein markers, as measured by the bead-based flow cytometry of enriched EVs (yellow field). The sizes of the circles represent the relative average normalized MFI signal for the indicated protein marker, while the biological variability (CV) is indicated by the shading of the circle borders. Cellular sources of specific circulating EV subsets are indicated by a solid line from cells to EVs. When multiple cell types contribute to EVs, the most frequent cellular source is depicted with a solid line, while less common or less probable sources are shown with dashed lines. We demonstrated a significant influence of sex (yellow icon), menopausal status (pink icon), age (violet icon) and smoking (blue icon) on inter-individual biological variability of specific circulating EV subsets and/or tetraspanin+ EVs. For more detailed information, please refer to the manuscript. Image created by Biorender.com.

Our results provide the first comprehensive, quantitative data towards the cell-origin atlas of the EV population in the blood of healthy humans. Imaging flow cytometry analysis of plasma detected median EV concentration across all 25 studied subsets of 2.3 x 10^7^/mL, which is in line with published flow cytometry studies on human circulating EVs (Johnsen et al. 2019; Bettin et al. 2022; Woud et al. 2022). Still, we observed up to a 221-fold variance in median concentrations among distinct EV subsets, with most numerous being EVs from activated platelets (PAC-1), followed by erythrocyte EVs (CD235a), activated granulocyte-derived EVs (CD66b), T lymphocyte EVs (CD152), EV released from APCs (HLA-DR), monocyte EVs (CD14), and activated endothelial EVs (CD90). Arraud et al. similarly observed with electron microscopy that 30% and 20% of circulating EVs were of platelet and erythrocyte origin, respectively (Arraud et al. 2014). In contrast, studies of the distribution of circulating EVs based on the RNA sequencing data underestimated the concentration of erythrocyte and neutrophil EVs and did not study endothelial EVs (Auber and Svenningsen 2022; Li et al. 2020). The abundance of the EV subsets in our study does not reflect the abundance of the parental cell types, as measured by the complete blood count in the donor blood. For example, erythrocytes were 22 times more numerous than platelets but showed 3 times lower EV concentration if comparing PAC-1+ to CD235a+ EV subsets. This likely reflects the low EV secretion rate for erythrocytes compared to other blood cells (Auber and Svenningsen 2022; Gamonet et al. 2020) and/or the higher secretion rate of EVs from activated platelets (Guerreiro et al. 2024).

Still, we have evidence of further complexity to circulating EV subset steady-state concentration regulation, since correlation analysis detected no strong patterns among EV subsets originating from the same cell type. If we focus on EVs originating from activated platelets, they were the most numerous circulating EVs when measuring PAC-1+ EVs (41.85 x 10^6^/mL), and the least numerous when measuring CD62P+ EVs (0.19 x 10^6^/mL). EV subsets carrying the general markers of platelets CD61 or CD41 had median concentrations of 2.03 x 10^6^/mL and 3.44 x 10^6^/mL, respectively. Large variability in platelet circulating EV subset levels could reflect the diversity in the biogenesis of platelet EVs, which can be released by plasma membrane budding, extrusion of multivesicular α-granules and cytoplasmic vacuoles, plasma membrane blistering, or “pearling” of platelets pseudopodia upon platelet activation (Heijnen et al. 1999; De Paoli et al. 2018). Further studies demonstrated that Ca^2+^-triggers release PS-exposing EVs from platelet plasma membrane, with lipid rafts playing an essential role (Wei, Malcor, and Harper 2018). Interestingly, upon activation, the PS-exposing subpopulation of platelets expressed higher levels of CD62P and lower levels of activated αIIbβ3 (PAC-1+) on their surface, compared to PS-negative platelets (Reddy et al. 2018), which could be replicated in the released EVs. PS exposure to the cell surface is recognised as one of the key ‘eat me’ signals for the clearance of apoptotic cells by macrophages (Segawa and Nagata 2015). In line with that, Matsumura et al. reported that circulating EVs not exposing PS on the membrane surface show slower clearance by macrophages in the mice blood (Matsumura et al. 2019). Accelerated clearance of PS+ EVs could thus explain the observed lower levels of the CD62P+ EV subset compared to the PAC1+ EV subset in our cohort. Interestingly, tetraspanin+ circulating EVs in healthy humans largely originated from platelets and showed a conserved pattern of protein marker expression across our study subjects, implying a similar mechanism of biogenesis in platelets. Among the three commonly used EV-related tetraspanins, only CD9 and CD63 seemed to be related to EVs of platelet origin, but not CD81. Altogether, release and clearance processes likely contribute to the observed differences in steady-state concentrations of EV subsets originating from the same cell type.

We have shown high inter-individual biological variability of circulating EV subset concentrations in healthy humans, with the granulocyte-derived CD66b+ EVs, CD81+ EVs, activated endothelial CD90+ EVs, and CD44+ EVs contributing most to the observed variability, according to PCA. The median CV across all EV subsets was 162%, with the highest and the lowest CV of 719% and 63% observed for activated endothelial CD90+ EV and CD9+ EV subsets, respectively. For easier visualisation, this translates to an almost 11,000-fold difference in CD90+ EV concentrations and only a 26-fold difference in CD9+EV concentrations among healthy individuals in our study. This variation surpasses even the high biological variation of commonly examined blood plasma analytes (Fest et al. 2018; Apoil et al. 2017) and underlines the necessity of establishing distinct reference intervals for each circulating EV subset intended for use as a biomarker, thus improving clinical decision-making. In comparison, tetraspanin+ circulating EVs exhibited lower variation in expression levels of cell-type specific protein markers (median CV of 60%) but are not fully representative of the whole EV population in blood, as they largely originate from platelets and to a lesser extent from lymphocytes. Of note, high inter-individual variability observed for some circulating EV subsets and expression of markers on tetraspanin+ EVs, including the CD90+ EVs and the CD4 marker, was due to high CVs observed in females (CV of 663% for CD90+ EVs and 127% for CD4 on tetraspanin+ EVs), supporting stratification of reference intervals by sex in such cases. Previous studies support the influence of female hormones on the blood CD90 concentrations and blood or endothelial cell surface expression of CD90 and CD4 (Bochev et al. 2022; Latorre et al. 2022; Elliott Williams et al. 2023). For example, increased concentrations of soluble CD90 are linked to endometriosis (Bochev et al. 2022) and the fluctuations in oestrogen and progesterone levels during the menstrual cycle and pregnancy influence the activity and regulation of CD4+ T cells (Polese et al. 2014). Interestingly, even EV subsets originating from the same cell type showed differing levels of variability. In the case of platelet EVs, CD41+, PAC-1+ and CD62P+ EV subsets had CVs of 138%, 153% and 637%, respectively. This could suggest the preferred use of CD41+ and PAC-1+ EVs as markers of platelets and activated platelets in blood as they are more consistent across healthy humans, but on the other hand, CD62P+ EV subsets might be richer in information regarding ongoing (pato)physiological processes. Further studies are needed to better understand the biological variability of circulating EVs and how it is affected by disease.

Here performed comprehensive association analysis supports that sex, menopausal status, age and smoking contribute to inter-individual biological variability observed in circulating EVs in healthy humans, but the influence was limited to specific EV subsets and/or tetraspanin+ EVs, as no global influence across circulating EVs was detected in PCA. This is supported by the studies in which age, sex and/or smoking status were shown to affect specific circulating EV subsets (Enjeti et al. 2017; Gustafson et al. 2015) and protein profiles on tetraspanin+ circulating EVs (Bæk, Varming, and Jørgensen 2016). In contrast, Eitan et al. implied the global effects of age on circulating EV concentration, but this might be due to the detection of all precipitated nanoparticles from blood, not just EVs, with NTA in their study (Eitan et al. 2017). Specifically, our study found higher concentrations of PAC-1+ platelet-derived EVs in females, indicating a sex-related pro-coagulant profile, consistent with reports of elevated platelet (CD62P, CD41, CD42a) -derived microvesicles in female (Gustafson et al. 2015). Furthermore, the pro-coagulative nature of microvesicles was higher during the luteal phase of the menstrual cycle, compared to the follicular phase (Toth et al. 2007), implicating the involvement of female sex hormones in this phenomenon. Here, smoking also influenced platelet-derived EVs, as smokers showed increased expression of CD41b and CD62P on tetraspanin+ circulating EVs, supporting previous findings by Mobarrez et al. that smoking even a single cigarette is reflected in elevated microparticles from platelets (Mobarrez et al. 2014). Additionally, smokers showed significantly lower levels of erythrocyte-derived CD235a+ circulating EVs in our study, aligning with findings of increased macrocytic erythrocytes in smokers, since smoking affects the morphology of the red blood cells (Aldosari et al. 2020). This could similarly cause perturbation of vesiculation from erythrocytes in our cohort. Interestingly, pre-menopausal females had significantly higher concentrations of leukocyte-derived CD100+, CD152+, and CD66b+ EVs than post-menopausal females. While immune variations occur in menopause, differences in immune cell-derived circulating EVs haven’t been reported previously. Others have shown decreases in microvesicles from erythrocytes (CD235a+) and endothelial cells (CD105+) in post-menopausal females (Gustafson et al. 2015; Enjeti et al. 2017), which was not reflected in our cohort. On the other hand, smoking reduced T lymphocytic CD8+ and monocytic CD14+ EVs in our study, supporting the modulatory effect of cigarette smoke on immune cells (Russell et al. 2022). Analysing endothelial cell-derived EVs, we observed a significant decrease in CD146+ EVs in early middle-aged adults compared to older adults, a novel finding as CD146’s role in ageing hasn’t been described before. We also noted a decrease in CD105+ EVs in early middle-aged adults compared to young adults, while Enjeti et al. reported a continuous decrease in CD105+ microvesicles across age groups (Enjeti et al. 2017).

Our study is the first to demonstrate high biological inter-individual variability in circulating EVs across healthy humans, which can be partly explained by the influence of sex, menopausal status, age and smoking on specific blood cells and endothelial cells. Still, several limitations of our study should be acknowledged. Our healthy human cohort was composed of Slovenian blood donors of South Slavic ethnicity, limiting exploration of the influence of race on circulating EVs. Furthermore, study subjects were not fasted, since eating a non-fatty meal before blood donation is a routine recommendation. However, statistical analysis found no influence of serum lipoprotein and insulin values on the study results. Although relying on self-reported data on smoking, exercise, height, and weight can introduce potential biases, we minimised those by performing in-person interviews conducted by the same medical doctor, rather than through self-administered questionnaires. Additionally, ours was a single time-point (cross-sectional) study, limiting our understanding of biological intra-individual variability of circulating EVs over time. Still, we employed strict inclusion criteria, so the study cohort represents apparently healthy adult population, and have performed comprehensive quality checks and reporting (MIBlood-EV, MIFlowCyte-EV) to ensure consistency and minimise the influence of pre-analytical variables and technical variability on the study outcomes.

In summary, our study is the first to comprehensively quantify blood EVs with respect to cell origin in a large cohort of healthy humans. It improves the understanding of the physiological circulating EV population, its biological variability and sources of observed variability. Our findings have important implications for fundamental research and clinical applications of blood EVs, especially as biomarkers of diverse diseases.

## Author contributions

Marija Holcar: Conceptualization; Formal analysis; Investigation; Methodology; Resources; Data curation; Visualization; Writing – original draft; Writing – review & editing.

Ivica Marić: Conceptualization; Funding acquisition; Investigation; Methodology; Resources; Data curation; Writing – review & editing.

Tobias Tertel: Formal analysis; Investigation; Methodology; Resources; Writing – review & editing.

Katja Goričar: Formal analysis; Investigation; Methodology; data curation; Writing – review & editing.

Urška Čegovnik Primožič: Formal analysis; Investigation; Methodology; Writing – review & editing.

Darko Černe: Methodology; Resources; Writing – review & editing.

Bernd Giebel: Methodology; Resources; Writing – review & editing.

Metka Lenassi: Conceptualization; Funding acquisition; Project administration; Resources; Supervision; Writing – original draft; Writing – review & editing.

## Supporting information

MIFlow-Cyt EV

MIBlood-EV plasma

MIBlood-EV serum

## Acknowledgements

The authors would like to thank Nina Mavec, for her help with the enrichment of EVs from blood plasma.

## Geolocation information

46.050980777270254, 14.518433998551663

