## Supplement file for "Comprehensive Phenotyping of Extracellular Vesicles in Blood of Healthy Humans – Insights into Cellular Origin and Biological Variability"

**Supplementary Material:**

**S1 Imaging flow cytometry analysis of circulating extracellular vesicles**

To phenotype circulating EVs with respect to their cell of origin, we used antibodies against 25 surface protein markers, listed in Supplementary Table 1 together with information on typical presence on blood or endothelial cells. Information on individual antibodies used in imaging flow cytometry is presented in Supplementary Table 2, while their allocation in specific assays is provided in Supplementary Table 3. Data on the calibration of imaging flow cytometry fluorescence signal to the molecules of equivalent soluble fluorochrome (MESF) signal is presented in Suppl. Figure 1. Laser settings and detection channels and filters used for imaging flow cytometry are presented in Supplementary Table 4, while the applied compensation matrix is depicted in Supplementary Table 5.

**Supplementary Table 1: Presence of surface protein markers, targeted in imaging flow cytometry, on blood or endothelial cells**

| | Leukocyte | | | | | | | | | |
|---|---|---|---|---|---|---|---|---|---|---|
| | Lymphocyte | | | Monocyte | | | | | | |
| | T-cell | B-cell | NK cell | Macrophage/monocyte | Dendritic cell | Granulocyte | Platelet | Erythrocyte | Endothelial cell | Stem/precursor cell |
| **CD90** | - | - | - | - | | - | - | - | + | + |
| CD71 | - | - | - | | - | | | - | + | + |
| **CD152** | + | + | - | - | - | - | - | | | |
| CD227 | + | + | | + | + | | | | | + |
| CD14 | - | - | - | + | | + | | | | |
| **CD66b** | - | - | - | | | + | - | - | - | |
| CD24 | - | + | - | + | - | + | - | - | - | - |
| CD100 | + | + | + | + | - | + | - | - | - | - |
| CD41 | - | - | - | - | - | - | + | - | - | + |
| **PAC-1** | | | | | | | + | | | |
| CD235a | - | - | - | - | - | - | | + | | + |
| **CD62P** | | | | | | | + | | + | |
| **HLA-DR** | + | + | | + | + | | | | | |
| CD81 | + | + | + | + | + | | - | - | + | + |
| CD13 | + | - | - | + | | + | - | - | + | + |
| CD206 | | | | + | + | | | | + | |
| HLA-ABC | + | + | + | + | + | + | + | - | + | |
| CD9 | + | + | | + | | + | + | | + | - |
| CD63 | + | + | + | + | - | + | + | | + | |
| CD29 | + | + | + | + | + | + | + | | + | + |

| | | | | | | | | | | |
|---|---|---|---|---|---|---|---|---|---|---|
| CD31 | **+** | **+** | **+** | **+** | | **+** | **+** | - | **+** | |
| CD61 | | | | **+** | | | **+** | | **+** | |
| CD82 | **+** | **+** | **+** | **+** | | **+** | **+** | - | **+** | **+** |
| **PS** | **+** | **+** | **+** | **+** | **+** | **+** | **+** | **+** | **+** | |
| CD44 | **+** | **+** | **+** | **+** | | **+** | - | **+** | **+** | |

PS – phosphatidylserine; Markers of cell activation are written in bold. The cellular origin of the markers is summarised after (“CD Marker Handbook Human and Mouse” 2016; Kalina et al. 2019; Grant et al. 2021; de Oliveira et al. 2023; Berckmans et al. 2019; Spurgeon and Frelinger 2022).

**Supplementary Table 2: Overview of antibodies used for phenotyping of circulating EVs with respect to their cell of origin.**

| **Antibody** | **Conjugate** | **Clone** | **Order Nr.** | **Company** | **Isotype** | **Volume used:** |
|---|---|---|---|---|---|---|
| **CD90** | APC | 5E10 | 1A-652-T100 | E | mouse IgG1, κ | 0.10 |
| **CD71** | PE | MEM-75 | 1P-235-T100 | E | mouse IgG1, κ | 0.25 |
| **CD152** | PE | BNI3 | 555853 | BD | mouse IgG2a, κ | 0.10 |
| **CD227** | FITC | HMPV | 559774 | BD | mouse IgG1, κ | 0.25 |
| **CD14** | APC | REA599 | 130-110-520 | M | human IgG1, rec | 0.10 |
| **CD66b** | APC | 6/40c | -* | LeuKoCom | mouse IgG1, κ | 0.10 |
| **CD24** | PE | M1/69 | 130-102-732 | M | rat IgG2bκ | 0.25 |
| **CD100** | PE | 133-1C6 | 1P-772-T100 | E | mouse IgM, κ | 0.25 |
| **CD41** | AF488 | MEM-06 | A4-309-T100 | E | mouse IgG1, κ | 0.25 |
| **PAC-1** | AF647 | PAC-1 | 362805 | BL | mouse IgM, κ | 0.10 |
| **CD235a** | APC | GA-R2 | 551336 | BD | mouse IgG2a, κ | 0.25 |
| **CD62P** | PE | AK-4 | 555524 | BD | mouse IgG1, κ | 0.10 |
| **HLA-DR** | ECD | Immu-357 | B92438 | BC | mouse IgG1, κ | 0.25 |
| **CD81** | FITC | JS64 | B25329 | BC | mouse IgG2a, κ | 0.25 |
| **CD13** | PE | QA19A12 | 111003 | BL | rat IgG2a, κ | 0.25 |
| **CD206** | BV421 | 15-2 | 321126 | BL | mouse IgG1, κ | 0.10 |
| **HLA-ABC** | FITC | B9.12.1 | IM1838U | BC | mouse IgG2a, κ | 0.25 |
| **CD9** | PE | MEM-61 | 1P-208-T100 | E | mouse IgG1, κ | 0.25 |
| **CD63** | APC | MEM-259 | 1A-343-T100 | E | mouse IgG1, κ | 0.25 |
| **CD29** | APC | REA1060 | 130-118-122 | M | human IgG1, rec | 0.10 |
| **CD31** | APC | REA730 | 130-110-670 | M | human IgG1, rec | 0.10 |
| **CD61** | FITC | SZ21 | IM1758 | BC | mouse IgG1, κ | 0.25 |
| **CD82** | PE | ASL-24 | 342104 | BL | mouse IgG1, κ | 0.25 |
| **PS** | AF488 | 1H6 | 16-256 | S | mouse IgG1, κ | 0.10 |
| **CD44** | APC | MEM-85 | 1A-221-T100 | E | mouse IgG2a, κ | 0.25 |

*kindly provided by LeuKoCom, Bernhard Singer, University Hospital Essen

The table lists the antibody, conjugated fluorophore, and the volume used for a 10 µL setup. BL - BioLegend; BD - BD Bioscience; BC - Beckman Coulter; M - Miltenyi Biotec; E - Exbio; S - Sigma-Aldrich; rec - recombinant

**Supplementary Table 3: Allocation of antibodies in imaging flow cytometry assays.**

| | Fluorophore 1 | Fluorophore 2 | Fluorophore 3 | Fluorophore 4 | Fluorophore 5 |
|---|---|---|---|---|---|
| **Tube 1** | CD9 | CD41 | - | CD29 | - |
| **Tube 2** | CD24 | PS | - | PAC-1 | - |
| **Tube 3** | CD13 | CD61 | - | CD31 | - |
| **Tube 4** | CD100 | HLA-ABC | - | CD90 | - |
| **Tube 5** | CD82 | - | HLA-DR | CD66b | - |
| **Tube 6** | CD152 | CD81 | - | CD44 | - |
| **Tube 7** | CD62P | CD227 | - | CD235a | - |
| **Tube 8** | CD71 | CD63 | - | CD14 | CD206 |

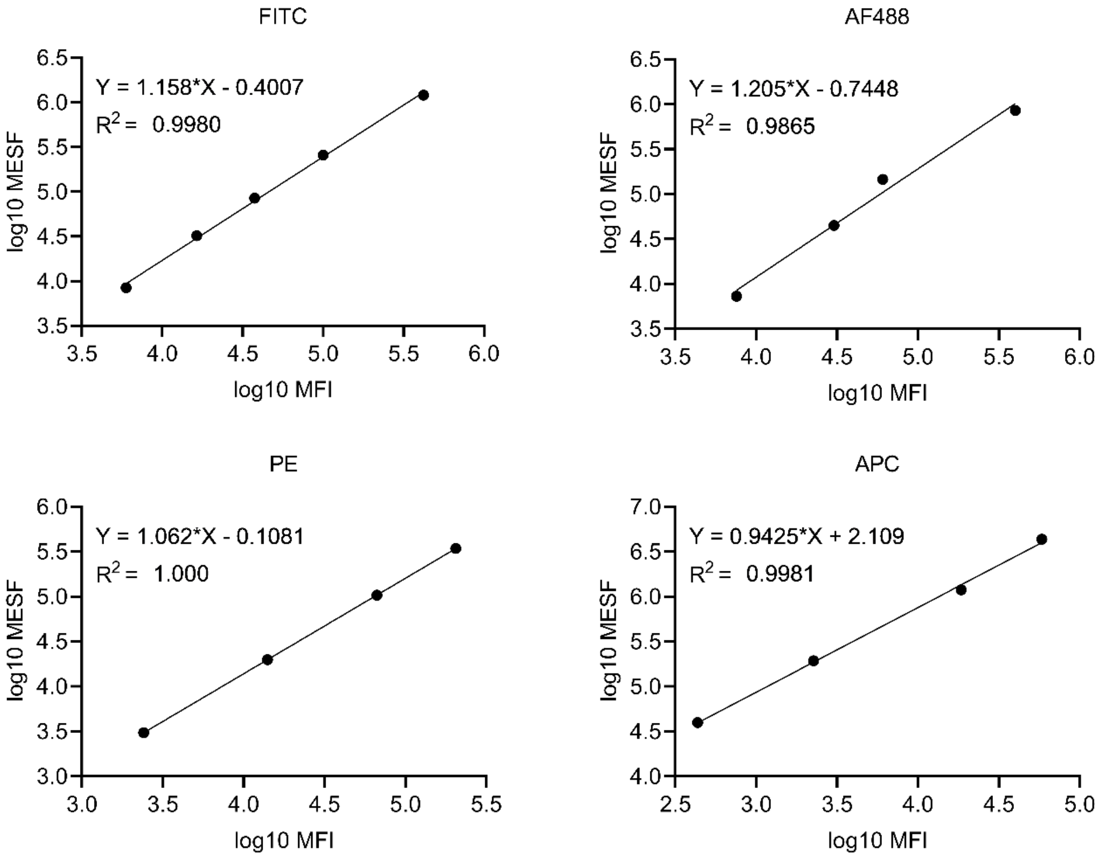


**Supplementary Figure 1: Calibration curve to convert fluorescence intensity readings into molecules of equivalent soluble fluorochrome of FITC, AF488, PE, and APC.**

The graph displays the linear regression analysis used to convert fluorescence intensity readings into molecules of equivalent soluble fluorochrome (MESF) units for each fluorophore. Data points represent the measured fluorescence intensity (y-axis) for standard beads with known MESF values (x-axis), with each fluorophore depicted on a separate graph.

**Supplementary Table 4: Laser settings and detection channels/filters used.**

| Laser [nm] | Used Power [mW] | Max. Power [mW] | Filter [nm] |
|---|---|---|---|
| 375 | 70 | 70 | BV421 (Ch07) 435-505 |
| 488 | 100 | 100 | FITC (Ch02) 480-560 |
| 561 | 200 | 200 | PE (Ch03) 560-595<br>ECD (Ch04) 595-642 |
| 648 | 150 | 150 | APC (Ch11) 642-745 |
| 785 (SSC) | 70 | 70 | SSC (Ch06) 756-780 |

**Supplementary Table 5. Applied compensation matrix for imaging flow cytometry of circulating EVs, stained with the antibodies.**

| | Ch1 | Ch2 | Ch3 | Ch4 | Ch5 | Ch6 | Ch7 | Ch8 | Ch9 | Ch10 | Ch11 | Ch12 |
|---|---|---|---|---|---|---|---|---|---|---|---|---|
| **Ch1** | 1 | 0.029 | 0.042 | 0 | 0 | 0 | 0 | 0 | 0 | 0 | 0 | 0 |
| **Ch2** | 0.051 | 1 | 0.05 | 0 | 0 | 0 | 0 | 0 | 0 | 0 | 0 | 0 |
| **Ch3** | 0 | 0.13 | 1 | 0 | 0 | 0 | 0 | 0 | 0.02 | 0 | 0 | 0 |
| **Ch4** | 0 | 0.064 | 0.49 | 1 | 0 | 0 | 0 | 0 | 0 | 0 | 0 | 0 |
| **Ch5** | 0 | 0.017 | 0.155 | 0 | 1 | 0 | 0 | 0 | 0 | 0 | 0 | 0 |
| **Ch6** | 0.015 | 0.02 | 0.04 | 0 | 0 | 1 | 0 | 0 | 0 | 0 | 0 | 0 |
| **Ch7** | 0.023 | 0.003 | 0.003 | 0 | 0 | 0 | 1 | 0 | 0.015 | 0 | 0 | 0 |
| **Ch8** | 0 | 0.032 | 0.008 | 0 | 0 | 0 | 0 | 1 | 0.012 | 0 | 0 | 0 |
| **Ch9** | 0 | 0.004 | 0.084 | 0 | 0 | 0 | 0 | 0 | 1 | 0 | 0 | 0 |
| **Ch10** | 0 | 0.002 | 0.041 | 0 | 0 | 0 | 0 | 0 | 0.084 | 1 | 0 | 0 |
| **Ch11** | 0 | 0.001 | 0.012 | 0 | 0 | 0 | 0 | 0 | 0.025 | 0 | 1 | 0 |
| **Ch12** | 0 | 0 | 0.003 | 0 | 0 | 0 | 0 | 0 | 0.013 | 0 | 0 | 1 |

### S2: Analysis of the correlation between serum lipoproteins and insulin levels and all measured EV characteristics

**Supplementary Table 6: Serum lipoprotein and insulin concentration correlation analysis**

| | | ApoA1 | ApoB | Insulin |
|---|---|---|---|---|
| Imaging flow cytometry | | | | |
| CD41 | Spearman's rho | 0.025 | 0.124 | 0.066 |
| | P | 0.726 | 0.081 | 0.395 |
| | N | 200 | 200 | 168 |
| PS | Spearman's rho | -0.014 | -0.124 | -0.117 |
| | P | 0.841 | 0.080 | 0.131 |
| | N | 200 | 200 | 168 |
| CD61 | Spearman's rho | -0.031 | 0.157 | -0.106 |
| | P | 0.660 | 0.026 | 0.173 |
| | N | 199 | 199 | 167 |
| HLA-ABC | Spearman's rho | -0.046 | -0.015 | -0.091 |
| | P | 0.514 | 0.833 | 0.242 |
| | N | 200 | 200 | 168 |
| CD81 | Spearman's rho | -0.083 | 0.091 | -0.050 |
| | P | 0.242 | 0.202 | 0.516 |
| | N | 200 | 200 | 168 |
| CD227 | Spearman's rho | -0.091 | -0.052 | -0.040 |
| | P | 0.199 | 0.465 | 0.605 |
| | N | 200 | 200 | 168 |
| CD63 | Spearman's rho | -0.045 | -0.023 | 0.056 |
| | P | 0.529 | 0.744 | 0.473 |
| | N | 200 | 200 | 168 |
| CD9 | Spearman's rho | -0.015 | 0.151 | 0.149 |
| | P | 0.834 | 0.033 | 0.055 |
| | N | 200 | 200 | 168 |
| CD24 | Spearman's rho | -0.197 | 0.061 | 0.066 |
| | P | 0.005 | 0.389 | 0.393 |
| | N | 200 | 200 | 168 |
| CD13 | Spearman's rho | -0.218 | 0.239 | 0.027 |
| | P | 0.002 | 0.001 | 0.728 |
| | N | 200 | 200 | 168 |
| CD100 | Spearman's rho | -0.154 | -0.203 | 0.004 |
| | P | 0.030 | 0.004 | 0.961 |
| | N | 200 | 200 | 168 |
| CD82 | Spearman's rho | -0.153 | 0.203 | 0.062 |
| | P | 0.031 | 0.004 | 0.423 |
| | N | 200 | 200 | 168 |
| CD152 | Spearman's rho | -0.050 | -0.114 | -0.035 |

| | | | | |
|---|---|---|---|---|
| | P | 0.482 | 0.107 | 0.652 |
| | N | 200 | 200 | 168 |
| CD62P | Spearman's rho | 0.040 | 0.043 | 0.006 |
| | P | 0.573 | 0.550 | 0.938 |
| | N | 200 | 200 | 168 |
| CD71 | Spearman's rho | -0.006 | 0.001 | 0.009 |
| | P | 0.938 | 0.989 | 0.906 |
| | N | 200 | 200 | 168 |
| HLA-DR | Spearman's rho | -0.085 | 0.158 | 0.223 |
| | P | 0.230 | 0.026 | 0.004 |
| | N | 200 | 200 | 168 |
| CD29 | Spearman's rho | -0.010 | 0.091 | 0.064 |
| | P | 0.891 | 0.202 | 0.407 |
| | N | 200 | 200 | 168 |
| PAC-1 | Spearman's rho | -0.096 | -0.179 | 0.027 |
| | P | 0.175 | 0.011 | 0.728 |
| | N | 200 | 200 | 168 |
| CD31 | Spearman's rho | -0.030 | 0.105 | -0.047 |
| | P | 0.673 | 0.141 | 0.542 |
| | N | 200 | 200 | 168 |
| CD90 | Spearman's rho | -0.082 | 0.000 | -0.010 |
| | P | 0.250 | 0.994 | 0.900 |
| | N | 200 | 200 | 168 |
| CD66b | Spearman's rho | -0.015 | -0.116 | -0.063 |
| | P | 0.832 | 0.101 | 0.418 |
| | N | 200 | 200 | 168 |
| CD44 | Spearman's rho | -0.155 | 0.122 | 0.094 |
| | P | 0.044 | 0.115 | 0.269 |
| | N | 169 | 169 | 141 |
| CD235a | Spearman's rho | -0.123 | 0.082 | -0.086 |
| | P | 0.083 | 0.247 | 0.268 |
| | N | 200 | 200 | 168 |
| CD14 | Spearman's rho | -0.020 | 0.083 | 0.041 |
| | P | 0.777 | 0.240 | 0.593 |
| | N | 200 | 200 | 168 |
| CD206 | Spearman's rho | -0.127 | 0.021 | 0.175 |
| | P | 0.076 | 0.766 | 0.024 |
| | N | 197 | 197 | 165 |
| Bead-based flow cytometry | | | | |
| CD3 | Spearman's rho | 0.126 | -0.043 | 0.064 |
| | P | 0.072 | 0.539 | 0.408 |
| | N | 206 | 206 | 170 |
| CD4 | Spearman's rho | 0.105 | -0.089 | -0.007 |

| | | | | |
|---|---|---|---|---|
| | P | 0.133 | 0.202 | 0.924 |
| | N | 206 | 206 | 170 |
| CD8 | Spearman's rho | 0.036 | -0.126 | 0.046 |
| | P | 0.606 | 0.071 | 0.554 |
| | N | 205 | 205 | 169 |
| HLA-DRDPDQ | Spearman's rho | -0.082 | -0.061 | 0.075 |
| | P | 0.239 | 0.385 | 0.332 |
| | N | 206 | 206 | 170 |
| CD105 | Spearman's rho | -0.098 | 0.026 | 0.033 |
| | P | 0.163 | 0.710 | 0.669 |
| | N | 206 | 206 | 170 |
| CD2 | Spearman's rho | 0.147 | -0.096 | 0.058 |
| | P | 0.034 | 0.168 | 0.450 |
| | N | 206 | 206 | 170 |
| CD1c | Spearman's rho | 0.045 | -0.173 | -0.017 |
| | P | 0.524 | 0.013 | 0.826 |
| | N | 204 | 204 | 169 |
| CD9 | Spearman's rho | 0.084 | 0.061 | -0.105 |
| | P | 0.228 | 0.381 | 0.174 |
| | N | 206 | 206 | 170 |
| HLA-ABC | Spearman's rho | 0.011 | -0.004 | -0.016 |
| | P | 0.872 | 0.956 | 0.833 |
| | N | 201 | 201 | 166 |
| CD63 | Spearman's rho | 0.147 | -0.002 | 0.086 |
| | P | 0.035 | 0.973 | 0.262 |
| | N | 206 | 206 | 170 |
| CD40 | Spearman's rho | -0.050 | 0.087 | 0.002 |
| | P | 0.480 | 0.217 | 0.982 |
| | N | 204 | 204 | 169 |
| CD62P | Spearman's rho | 0.131 | 0.022 | 0.074 |
| | P | 0.060 | 0.750 | 0.336 |
| | N | 206 | 206 | 170 |
| CD81 | Spearman's rho | -0.108 | -0.003 | -0.016 |
| | P | 0.123 | 0.966 | 0.840 |
| | N | 206 | 206 | 170 |
| CD146 | Spearman's rho | -0.048 | -0.055 | 0.020 |
| | P | 0.490 | 0.433 | 0.794 |
| | N | 206 | 206 | 170 |
| CD41b | Spearman's rho | 0.105 | 0.018 | 0.034 |
| | P | 0.133 | 0.794 | 0.657 |
| | N | 206 | 206 | 170 |
| CD42a | Spearman's rho | -0.019 | 0.002 | 0.018 |
| | P | 0.782 | 0.978 | 0.817 |

| | | | | |
|---|---|---|---|---|
| | N | 206 | 206 | 170 |
| CD24 | Spearman's rho | 0.067 | -0.106 | -0.015 |
| | P | 0.340 | 0.129 | 0.848 |
| | N | 206 | 206 | 170 |
| CD44 | Spearman's rho | -0.018 | -0.067 | 0.016 |
| | P | 0.801 | 0.341 | 0.837 |
| | N | 206 | 206 | 170 |
| CD29 | Spearman's rho | -0.042 | 0.062 | 0.031 |
| | P | 0.547 | 0.379 | 0.690 |
| | N | 206 | 206 | 170 |
| CD69 | Spearman's rho | -0.065 | -0.038 | 0.094 |
| | P | 0.357 | 0.589 | 0.222 |
| | N | 206 | 206 | 170 |
| CD45 | Spearman's rho | 0.079 | -0.070 | 0.104 |
| | P | 0.260 | 0.321 | 0.176 |
| | N | 206 | 206 | 170 |
| CD31 | Spearman's rho | -0.036 | 0.003 | -0.031 |
| | P | 0.610 | 0.965 | 0.691 |
| | N | 205 | 205 | 170 |
| CD14 | Spearman's rho | 0.083 | -0.046 | 0.002 |
| | P | 0.236 | 0.510 | 0.983 |
| | N | 206 | 206 | 170 |
| Nanoparticle tracking analysis | | | | |
| EV concentration | Spearman's rho | 0.029 | 0.056 | -0.017 |
| | P | 0.676 | 0.421 | 0.827 |
| | N | 208 | 208 | 174 |
| Modal diameter | Spearman's rho | 0.087 | -0.076 | -0.021 |
| | P | 0.214 | 0.277 | 0.787 |
| | N | 208 | 208 | 174 |

None of the analysed correlations were significant considering the Bonferroni correction for the number of comparisons.

### S3: Whole blood analysis of the study subjects

**Supplementary Table 7: Whole blood analysis – complete blood count+**

| Characteristic | Unit | Median (25-75%) | Reference values* |
|---|---|---|---|
| Leukocytes | x $10^6$/mL | 6.22 (5.42-7.16) | 4-10 |
| Neutrophils | x $10^6$/mL | 3.48 (2.86-4.15) | 1.5-7.4 |
| | % of leuko | 56 (50.58-61.03) | 0.4-0.8 |
| Lymphocytes | x$10^6$/mL | 2.02 (1.63-2.39) | 1.1-3.5 |
| | % of leuko | 32.7 (27.48-36.55) | 0.2-0.4 |
| Monocytes | x $10^6$/mL | 0.51 (0.42-0.59) | 0.2-0.92 |
| | % of leuko | 7.99 (6.89-9.33) | 0.02-0.1 |
| Eosinophils | x $10^6$/mL | 0.15 (0.10-0.23) | 0.02-0.67 |
| | % of leuko | 2.33 (1.62-3.57) | 0.01-0.06 |
| Basophils | x $10^6$/mL | 0.065 (0.049-0.084) | 0.0-0.13 |
| | % of leuko | 1.05 (0.82-1.28) | 0.-0.02 |
| Erythrocytes | x $10^9$/mL | 4.91 (4.63-5.28) | 3.8-5.5 |
| Haemoglobin | mg/mL | 143 (133.3-152) | 120-170 |
| Haematocrit | mL/mL | 0.44 (0.41-0.47) | 0.36-0.5 |
| Platelets | x $10^6$/mL | 221 (191-249.5) | 150-410 |

+ Routine laboratory analysis, *Slovenian haematological guidelines (Zver et al. 2018)

### S3: Analysis of the quality of plasma and purity of enriched EV samples used in the study

To understand the contribution of typical confounders (lipoproteins, non-physiological platelet-derived EVs, haemolysis) when studying circulating EVs, we assessed the quality of plasma used in the study and evaluated the purity of EV samples enriched from those plasma samples. A summary of our quality assessment is also presented as MIBlood-EV reports attached to the manuscript.

The complete blood count of plasma samples on a routine haematology analyser showed undetectable levels of erythrocytes and platelets (Supplementary Table 7). To evaluate if the remaining (although undetectable) erythrocytes in plasma or their lysis contribute to non-physiological erythrocyte-derived EVs and other study outcomes, we measured erythrocytes (in whole blood and plasma), CD235a+ circulating EV concentrations, and free haemoglobin and performed correlation analysis. No correlation was found for any of the measured analytes, except haemoglobin concentration in the whole blood correlated with free haemoglobin concentration in the plasma ($\rho = 0.3354$, $p = 0.00000073$), which further supported the quality of plasma preparation.

To evaluate if the remaining (although undetectable) platelets in plasma contribute to non-physiological platelet-derived EVs and other study outcomes, we measured platelets (in whole blood and plasma), platelet-derived circulating EVs, and plasma PF4 as a marker of platelet activation, and performed correlation analysis (Supplementary Figure 2). No correlation was found for any of the measured analytes, except platelet counts in the whole blood correlated with plasma PF4 levels ($\rho = 0.3020$, $p = 0.000009$, Supplementary Figure 2a), CD62P+ tetraspanin+ circulating EVs and CD63+ tetraspanin+ circulating EVs ($\rho = 0.3028$, $p = 0.0000096$ and $\rho = 0.3148$, $p = 0.0000041$, Supplementary Figure 2b and c), which further supports absence of major platelet activation during plasma preparation and the physiological nature of circulating platelet-derived- EVs.

Finally, we measured lipoproteins ApoA1 and ApoB in EV samples and showed our method of EV enrichment from plasma removed a substantial portion of lipoproteins, as only 0.0057% of ApoA1 and 0.0017% of ApoB (92 out of 208 EV samples were below the limit of detection) were still detected (Supplementary Table 8, Supplementary Figure 2d). Importantly, no correlation was found for any of the measured analytes with any of the study outcomes, except ApoB concentration in EV-enriched samples correlated with serum ApoB levels ($\rho = 0.4426$, $p = 0.0000007$), which further supports the consistency of EV-enrichment process across study subjects.

**Supplementary Table 8: Additional analysis of plasma**

| Characteristic | Unit | Median (25-75%) |
|---|---|---|
| Erythrocytes | x $10^9$/mL | 0.00 (0.00-0.00) |
| Haemoglobin | mg/mL | 0.00 (0.00-0.00) |
| Platelets | x $10^6$/mL | 0.00 (0.00-0.00) |
| Platelet factor 4 | ng/mL | 1469 (781-2370) |
| Free haemoglobin | mg/mL | 0.010 (0.006-0.016) |

**Supplementary Table 9: Analysis of lipoprotein concentrations in enriched circulating EV**

| Characteristic | Unit / Category | Median (25-75%) | N (%) |
|---|---|---|---|
| ApoA1 | ng/mL | 90.5 (70.5-125.1) | |

| | | | |
|---|---|---|---|
| ApoB100* | <8.5 ng/mL | | 92 (44.2) |
| | ng/mL / >8.5 ng/mL | 15.6 (12.3-20.7) | 116 (55.8) |

ApoA1 – apolipoprotein A1; ApoB100 – apolipoprotein B100; *in 92 samples values were below the limit of detection (<8.5 ng/mL), only subjects with values larger than 8.5 ng/mL were included in the calculation of the median.

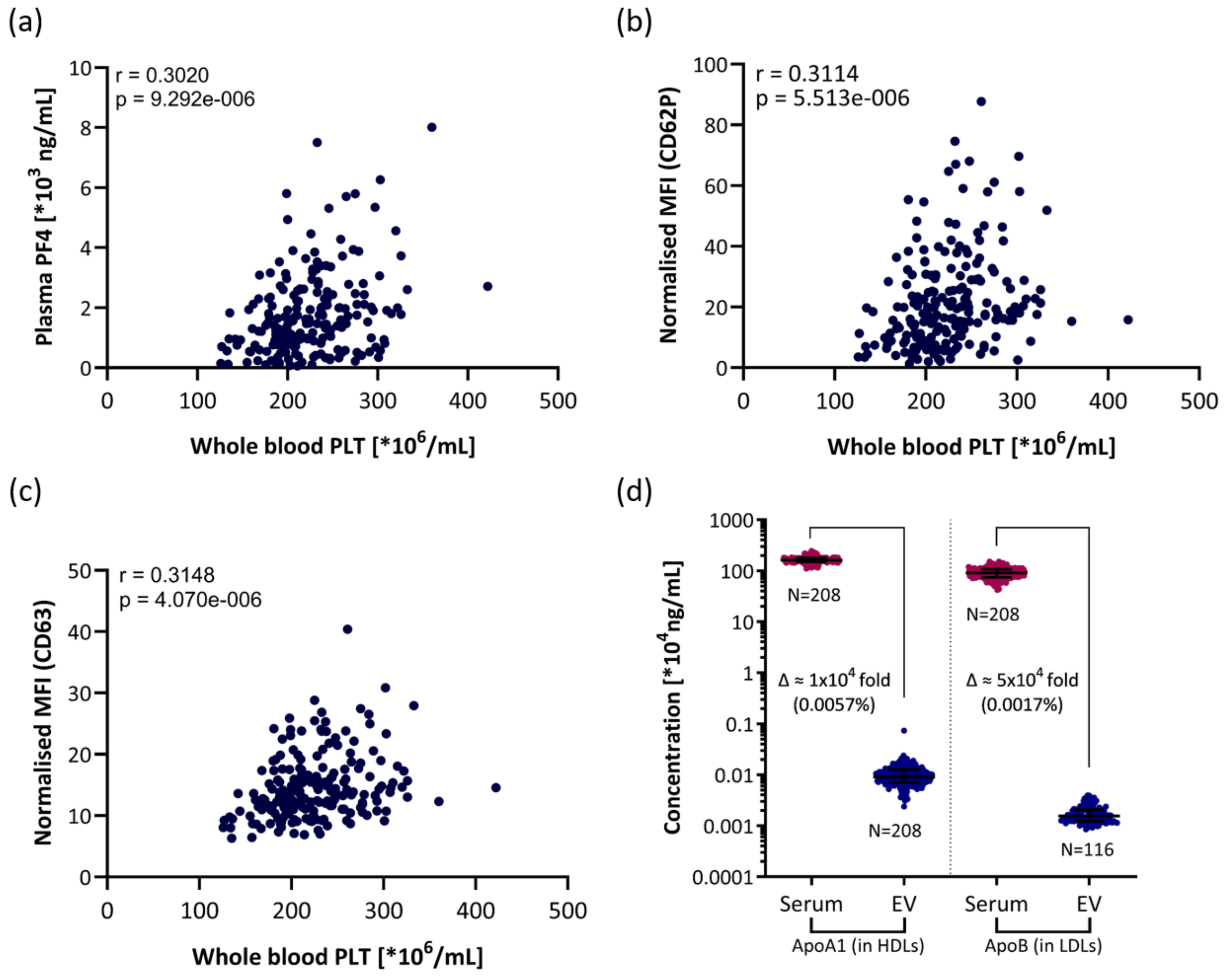


**Supplementary Figure 2: Analysis of the quality of plasma and purity of enriched EV samples used in the study.**

**(a)** Correlation between concentration of platelet factor 4 in plasma and concentration of platelets in whole blood. Correlation between normalised MFI of CD62+ **(b)** and CD63+ **(c)** tetraspanin+ circulating EVs and concertation of platelets in the whole blood. Each dot represents one study subject. **(d)** Apolipoproteins concentrations in serum and in EV samples enriched from plasma, reported as median with interquartile range. The concentration of ApoB in 92 samples of EVs after the enrichment was below the limit of detection (<8.5 $x10^{-6}$ mg/mL) and is not represented on the graph. ApoA1 – apolipoprotein A1; ApoB – apolipoprotein B; HDL – high-density lipoprotein; LDL – low-density lipoprotein; PF4 – platelet factor 4; PLT – platelets.

**S4 Nanoparticle Tracking Analysis of EV samples enriched from the blood plasma of healthy adults**

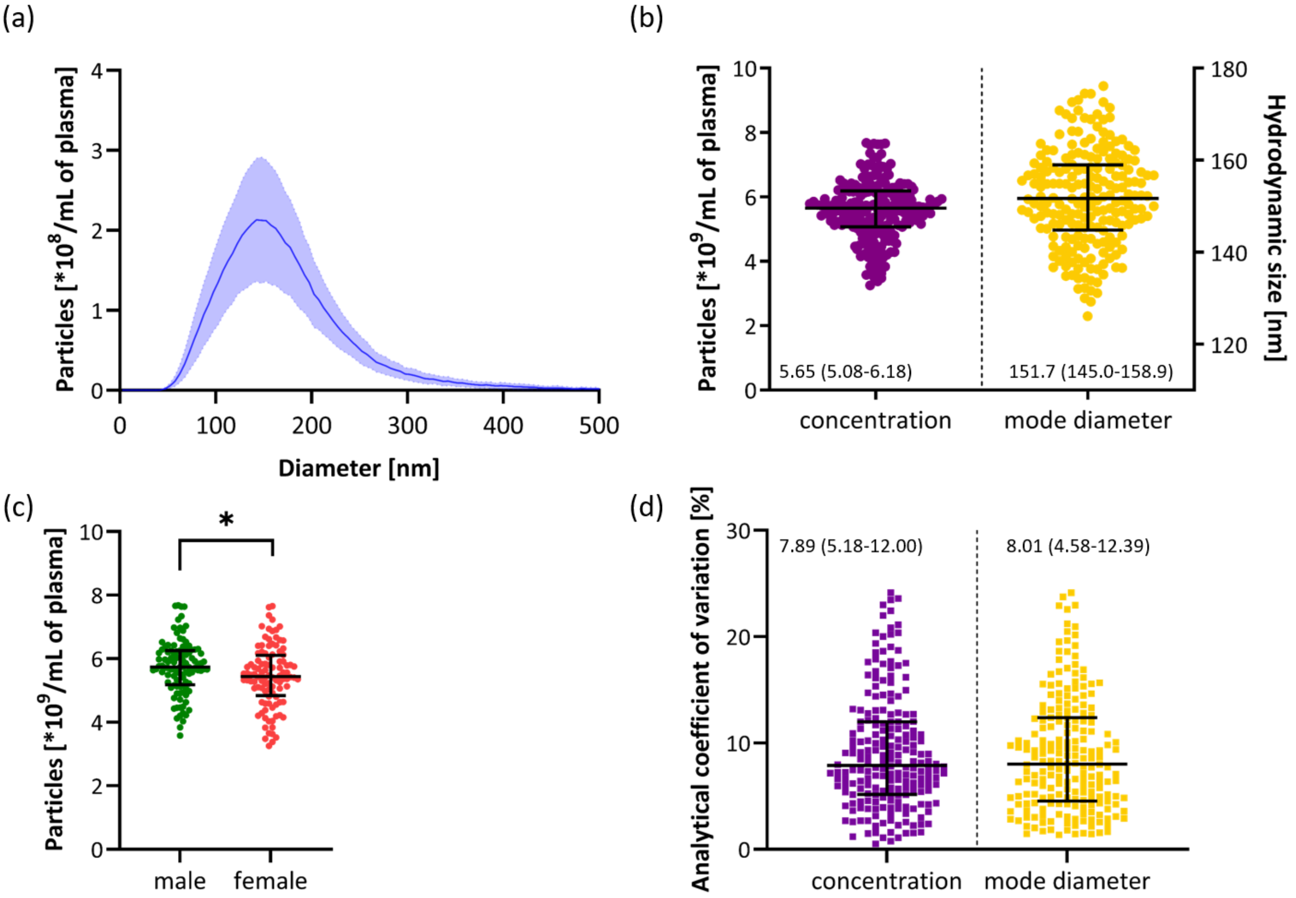


**Supplementary Figure 3: Results of nanoparticle tracking analysis of EV samples after enrichment of from blood plasma of healthy adults.**

**(a)** Mean of all measurements (line) with standard deviation (shaded area). **(b)** Concentration and mode diameter of particles in enriched circulating EV samples numerically represented as median (25-75%). Each dot represents one study subject. **(c)** Influence of sex on the concentration of measured particles. Each dot represents one study subject. Lines present the median with an interquartile range. * - $p < 0.05$. **(d)** Analytical coefficient of variation of particle concentration and mode diameter measurements numerically represented as median (25-75%). Each dot represents one study subject.

**S5 Multiplex bead-based flow cytometry analysis of tetraspanin+ circulating EVs**

To phenotype tetraspanin+ circulating EVs with respect to their cell of origin, we used antibodies against 23 surface protein markers, listed in Supplementary Table 9 together with information on typical presence on blood or endothelial cells.

**Supplementary Table 10: Presence of surface markers, included in multiplex bead-based tetraspanin+ circulating EV analysis, on various cell types**

| | Leukocyte | | | | | | Platelet | Erythrocyte | Endothelial cell | Stem/precursor cell |
|---|---|---|---|---|---|---|---|---|---|---|
| | Lymphocyte: T-cell | Lymphocyte: B-cell | Lymphocyte: NK cell | Monocyte: Macrophage/monocyte | Monocyte: Dendritic cell | Granulocyte | | | | |
| CD2 | + | + | + | | | - | - | - | | |
| CD3 | + | - | - | - | | - | - | - | - | - |
| CD8 | + | - | + | - | | - | - | - | - | - |
| CD1c | + | + | - | + | + | - | - | - | | |
| CD4 | + | - | - | + | | - | - | - | - | - |
| CD14 | - | - | - | + | | + | | | | |
| CD24 | - | + | - | - | - | + | - | - | - | - |
| CD45 | + | + | + | + | + | + | - | - | - | + |
| CD41b | - | - | - | - | - | - | + | - | - | + |
| CD42a | - | - | - | - | - | - | + | - | - | + |
| **CD62P** | | | | | | | + | | + | |
| **HLA-DRDPDQ** | + | + | | + | + | | | | | |
| CD40 | - | + | - | + | + | - | | - | + | + |
| CD81 | + | + | + | + | + | | - | - | + | + |
| CD105 | - | - | - | + | | - | - | - | + | + |
| CD146 | + | - | - | - | | - | - | - | + | |
| CD69 | + | + | + | + | | + | + | | | |
| HLA-ABC | + | + | + | + | + | + | + | - | + | |
| CD9 | + | + | | + | | + | + | | + | - |
| CD63 | + | + | + | + | - | + | + | | + | |
| CD29 | + | + | + | + | + | + | + | | + | + |
| CD31 | + | + | + | + | | + | + | - | + | |
| CD44 | + | + | + | + | | + | - | + | + | |

Markers of cell activation are written in bold. The cellular origin of the markers is summarised after ("CD Marker Handbook Human and Mouse" 2016; Kalina et al. 2019; Grant et al. 2021; de Oliveira et al. 2023; Berckmans et al. 2019; Spurgeon and Frelinger 2022).

#### S6 Tetraspanin+ circulating EVs reveal distinct profiles when compared to the overall circulating EV population.

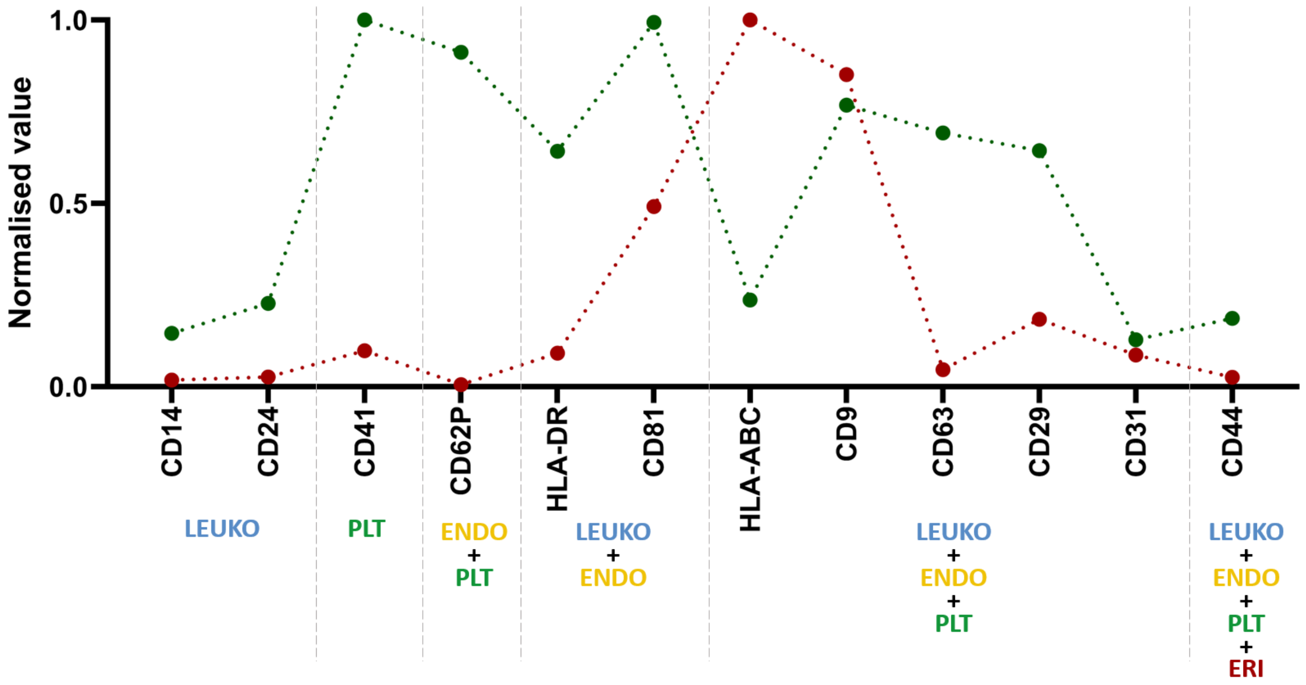


**Supplementary Figure 4: Tetraspanin+ circulating EVs reveal distinct profiles when compared to the overall circulating EV population.**

Red dots and line represent the circulating EV subset concentrations normalised to the CD41+ EV subset concentration (the highest among included EV subset concentrations), while green dots and line represent the expression levels of markers on tetraspanin+ circulating EVs normalised to the HLA-ABC MFI signal (the highest among included surface markers). Of note, we used antibodies against CD41 and HLA-DR in imaging flow cytometry, and antibodies against CD41b and HLA-DRDPDQ in multiplex bead-based flow cytometry, respectively. ENDO – endothelial cells; LEUKO – Leukocytes; PLT – platelets; ERI – erythrocytes.

**S7 Comparison of variability in the concentrations of circulating EV subsets or marker expression levels on tetraspanin+ circulating EVs between males and females**

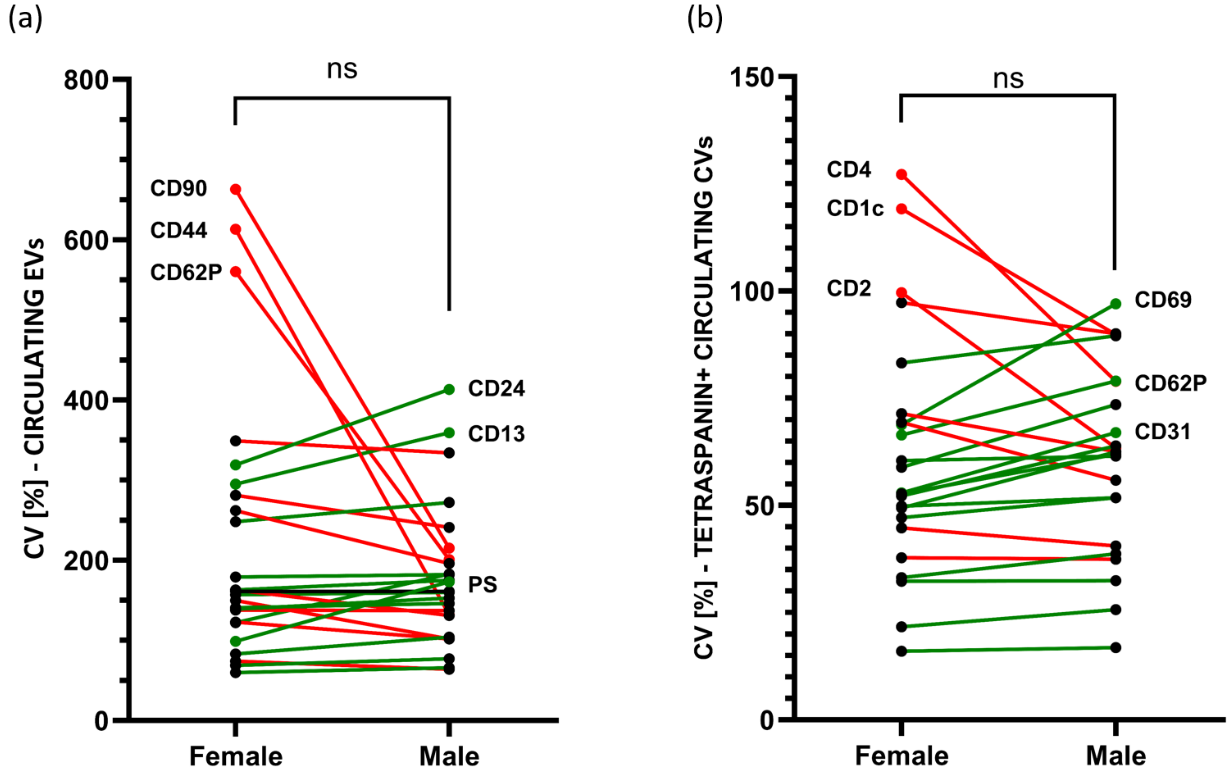


**Supplementary Figure 5: variability in the concentrations of circulating EV subsets or marker expression levels on tetraspanin+ circulating EVs between males and females. (a)** Comparison of coefficients of variations (CV) for measured concentrations of circulating EV subsets between females and males. **(b)** Comparison of CVs for marker expression levels on tetraspanin+ circulating EVs between females and males. Dots represent medians of measured values, with those showing the highest difference between sexes named. Red line - %CV was higher in females, green line - %CV was higher in males; ns – nonsignificant (Mann-Whitney test).
