## Supplementary material for "Comprehensive Phenotyping of Extracellular Vesicles in Blood of Healthy Humans – Insights into Cellular Origin and Biological Variability": MIFlow-Cyt EV

| **Framework Criteria** | **What to report** | **Please complete each criterion** |
|---|---|---|
| 1.1 Preanalytical variables conforming to MISEV guidelines. | Preanalytical variables relating to EV sample including source, collection, isolation, storage, and any others relevant and available in the performed study. | M&M |
| 1.2 Experimental design according to MIFlowCyt guidelines. | EV-FC manuscripts should provide a brief description of the experimental aim, keywords, and variables for the performed FC experiment(s) using MIFlowCyt checklist criteria: 1.1, 1.2, and 1.3, respectively. Template found at www.evflowcytometry.org. | M&M |
| 2.1 Sample staining details | State any steps relating to the staining of samples. Along with the method used for staining, provide relevant reagent descriptions as listed in MIFlowCyt guidelines (Section 2.4 Fluorescence Reagent(s) Descriptions). | M&M |
| 2.2 Sample washing details | State any steps relating to the washing of samples. | Not required |
| 2.3 Sample dilution details | All methods and steps relating to sample dilution. | M&M |
| 3.1 Buffer alone controls. | State whether a buffer-only control was analyzed at the same settings and during the same experiment as the samples of interest. If utilized it is recommended that all samples be recorded for a consistent set period of time e.g. 5 minutes, rather than stopping analysis at a set recorded event count e.g. 100,000 events. This allows comparisons of total particle counts between controls and samples. | M&M |
| 3.2 Buffer with reagent controls. | State whether a buffer with reagent control was analyzed at the same settings, same concentrations, and during the same experiment as the samples of interest. If used state what the results were. | M&M |
| 3.3 Unstained controls. | State whether unstained control samples were analyzed at the same settings and during the same experiment as stained samples. If used, state what the results were, preferably in standard units. | M&M |

| | | |
|---|---|---|
| 3.4 Isotype controls. | The use of isotype controls is applicable to immunofluorescence labelling only. State whether isotype controls were analyzed at the same settings and during the same experiment as stained samples. If utilized, state which antibody they are matched to, the concentration used, and what the results were (Section 4.2, 4.3, 4.4). Due to conjugation differences between manufacturers if should be stated if the isotype controls are from the same manufacturer as the matched antibodies. | M&M |
| 3.5 Single-stained controls. | State whether single-stained controls were included. If used state whether the single-stained controls were recorded using the same settings, dilutions, and during the same experiment as stained samples and state what the results were, preferably in standard units (Section 4.2, 4.3, 4.4). | M&M, Single stained samples were used only in establishing the protocol to identify any interferences. |
| 3.6 Procedural controls. | State whether procedural controls were included. If used, state the procedure and if the procedural controls were acquired at the same settings and during the same experiment as stained samples. | Not required |
| 3.7 Serial dilutions. | State whether serial dilutions were performed on samples and note the dilution range and manner of testing. The fluorescence and/or scatter signal intensity would ideally be reported in standard units (see Section 4.3, 4.4) but arbitrary units can also be used. This data is best reported by plotting the recorded number events/concentration over a set period of time at different sample dilution. The median fluorescence intensity at each of the dilutions should also ideally be plotted on the same or a separate plot. | Based on previous publications. |
| 3.8. Detergent treated EV-samples | State whether samples were detergent treated to assess lability. If utilized, state what detergent was used, the end concentration of the detergent, and what the results were of the lysis. | M&M |
| 4.1 Trigger Channel(s) and Threshold(s). | The trigger channel(s) and threshold(s) used for event detection. Preferably, the fluorescence calibration (Section 4.3) and/or scatter calibration (Section 4.4) should be used in order to report the trigger channel(s) and threshold(s) in standardized units. | M&M, Supplemental Tables. Threshold is automatically set from ISX (all channel are used to calculate the treshold) |
| 4.2 Flow Rate / Volumetric quantification. | State if the flow rate was quantified/validated and if so, report the result and how they were obtained. | M&M |

| | | |
|---|---|---|
| 4.3 Fluorescence Calibration. | State whether fluorescence calibration was implemented, and if so, report the materials and methods used, catalogue numbers, lot numbers, and supplied reference units for the standards. Fluorescence parameters may be reported in standardized units of MESF, ERF, or ABC beads. The type of regression used, and the resulting scatter plot of arbitrary data vs standard data for the reference particles should be supplied. | The system has been calibrated with MESF beads. ECD and BV421 are not calibrated (not available). AF647 was not available at the time of the experiment. |
| 4.4 Light Scatter Calibration. | State whether and how light scatter calibration was implemented. Light scatter parameters may be reported in standardized units of nm2, along with information required to reproduce the model. | The device is limited in detecting the scatterer, which is why no statement can be made about the size (See Görgens et al 2019., JEV). |
| 5.1 EV diameter/surface area/volume approximation. | State whether and how EV diameter, surface area, and/or volume has been calculated using FC measurements. | Not performed |
| 5.2 EV refractive index approximation. | State whether the EV refractive index has been approximated and how this was done. | Not performed |
| 5.3 EV epitope number approximation. | State whether EV epitope number has been approximated, and if so, how it was approximated. | Not performed |
| 6.1 Completion of MIFlowCyt checklist. | Complete MIFlowCyt checklist criteria 1 to 4 using the MIFlowCyt guidelines. Template found at www.evflowcytometry.org. | See above for 1-3. 4: Compensation matrix is included in the supplement tables. Gating strategy based on previous publication, mentioned in the M&M |

| | | |
|---|---|---|
| 6.2 Calibrated channel detection range | If fluorescence or scatter calibration has been carried out, authors should state whether the upper and lower limits of a calibrated detection channel were calculated in standardized units. This can be done by converting the arbitrary unit scale to a calibrated scaled, as discussed in Section 4.3 and 4.4, and providing the highest unit on this scale and the lowest detectable unit above the unstained population. The lowest unit at which a population is deemed ‘positive’ can be determined a variety of ways, including reporting the 99th percentile measurement unit of the unstained population for fluorescence. The chosen method for determining at what unit an event was deemed positive should be clearly outlined. | Gates were adjusted manually. |
| 6.3 EV number/concentration. | State whether EV number/concentration has been reported. If calculated, it is preferable to report EV number/concentration in a standardized manner, stating the number/concentration between a set detection range. | We diluted the samples appropriately to reach the optimal detection range of the system |
| 6.4 EV brightness. | When applicable, state the method by which the brightness of EVs is reported in standardized units of scatter and/or fluorescence. | Not performed |
| 7.1. Sharing of data to a public repository. | Provide a link to the experimental data in a public data repository. | On demand |
