## Supplementary material for "Comprehensive Phenotyping of Extracellular Vesicles in Blood of Healthy Humans – Insights into Cellular Origin and Biological Variability": MIBlood-EV plasma

INTERNATIONAL SOCIETY for EXTRACELLULAR VESICLES

### MIBlood-EV

Standardized Reporting Tool for Blood EV Research (Human)

#### STUDY INFORMATION

| | | | |
|---|---|---|---|
| 1.0 **Manuscript title** | Comprehensive Phenotyping of Extracellular Vesicles in Blood of Healthy Humans – Insights into Cellular Origin and Biological Variability | | |
| 1.1 **Corresponding author (Name and Email)** | Metka Lenassi, | | |
| 1.2 **Institution name** | Institute of Biochemistry and Molecular Genetics, Faculty of Medicine, University of Ljubljana, Ljubljana, Slovenia; EU | | |
| 1.3 **Time period of experiment (e.g. 2022-2024)** | 7.2020 - 4.2023 | 1.4 **Number of samples** | 208 |
| 1.5 **Cargo of interest** | ☑ Vesicles ☑ Protein ☐ RNA ☐ DNA ☐ Other: lipoproteins | | |
| 1.6 **Biospecimen type** | ☑ Plasma ☐ Serum | 1.7 **Biospecimen state** | Frozen |
| 1.8 **Source of frozen specimens** | Prepared by the lab | 1.9 **Years of collection (range)** | 7.2020 - 11.2020 |

#### BLOOD COLLECTION AND PROCESSING

| | | | |
|---|---|---|---|
| 2.0 **Patient fasting status** | Non-Fasted | 2.1 **Fasting length (e.g. hours/days)** | / |
| 2.2 **Anatomical access site** | cubital vein | 2.3 **Needle diameter (e.g. gauge)** | 16G |
| 2.4 **Blood volume collected (mL)** | 7.5 | | |
| 2.5 **Plasma anticoagulant** | ☑ EDTA ☐ Citrate ☐ Heparin ☐ Other: | | |
| 2.6 **Serum tube type** | | 2.7 **Serum clotting time (minutes)** | |
| 2.8 **Time between collection and first centrifugation (range in hours)** | 0-2.5 | | |
| 2.9 **Transport temperature** | Room temp (~22°C) | 2.10 **Transport condition of tubes** | Vertical |
| 2.11 **Centrifuge brand and model** | Eppendorf Centrifuge 5702 | | |
| 2.12 **Bucket rotor type** | Swing | 2.13 **Number of centrifugation cycles** | Two |
| 2.14a **1st Centrifugation speed (RCF in x g)** | 2500 | 2.14b **1st Centrifugation time (minutes)** | 15 |
| 2.15 **1st Rotor brake** | Off | 2.16 **1st Centrifugation temperature** | Room temp (~22°C) |
| 2.17a **2nd Centrifugation speed (RCF in x g)** | 2500 | 2.17b **2nd Centrifugation time (minutes)** | 15 |
| 2.18 **2nd Rotor brake** | Off | 2.19 **2nd Centrifugation temperature** | Room temp (~22°C) |
| 2.20 **Additional processing steps (e.g. filtration)** | / | | |
| 2.21 **Storage tubes (brand, type, source, catalog number)** | Greiner bio-one, Cryo.s Freezing Tubes, 2 ml, #122277 | | |
| 2.22 **Storage temperature** | -80°C | 2.23 **Length of storage (range in years)** | 2-3 |

#### PLASMA/SERUM QUALITY CONTROL

| | | | |
|---|---|---|---|
| 3.0 **Number of freeze-thaw cycles (range)** | 1-2 | | |
| 3.1 **Thawing temperature** | on ice | 3.2 **Thawing duration (minutes)** | 120 |

##### Hemolysis

| | | | |
|---|---|---|---|
| 3.3 **Presence of hemolysis** | Yes | 3.4 **Frequency of hemolyzed samples (e.g. <25%, 25-50%)** | <0.5% |
| 3.5 **Method used** | Spectrophotometry | 3.6 **RBC count (Median, 95% CI, N)** | 0; (0.0007+/-0.0003)x10^6/mL; 208 |
| 3.7 **RBC counter brand and type** | Abbot, CELL-DYN Ruby Hematology ANalyzer | | |
| 3.8 **Spectrophotometry hemoglobin concentration (mean g/L)** | 0-0.102 (min-max) | | |
| 3.9 **Spectrophotometer brand, model and wavelength measured (e.g. 414 nm)** | Biotek, Synergy 2, A415, A380, A450 (Harboe method) | | |
| 3.10 **Hemolized samples were discarded** | No | | |

#### Platelets

| | | | |
|---|---|---|---|
| 3.11 Presence of platelets | Yes | 3.12 Method used (e.g. Flow Cytometry) | Hematological Analyzer |
| 3.13 Marker(s) used (e.g. CD61, CD41) | / | | |
| 3.14 Concentration (median, 95% CI, N) | 0; (0.1318+/-0.0451)x10^6/mL; 208 | | |
| 3.15 Platelet counter instrument brand, type and limit of detection (cells/L) | Abbott, Cell-Dyn Ruby Hematology Analyzer; limit of detection not known | | |
| 3.16 Flow cytometer brand and type | / | | |
| 3.17 Flow cytometry size and fluorescence ranges of detection in nanometers and MESF, respectively | / | | |

#### Lipoproteins

| | | | |
|---|---|---|---|
| 3.18 Presence of lipoproteins | Yes | 3.19 Method used (WB, ELISA, FC) | Other: ELISA |
| 3.20 Spectrophotometry L-index | / | | |
| 3.21 Spectrophotometer brand, model and wavelength measured (e.g. 700 nm) | / | | |
| 3.22 WB Marker(s) used (e.g. Apo B) | / | | |
| 3.23 Western blot images provided in manuscript? | | | |
| 3.24 Flow cytometry marker(s) used (e.g. ApoB) | / | | |
| 3.25 Flow cytometry concentration (median, 95% CI, N) | / | | |
| 3.26 Flow cytometer brand and type | / | | |
| 3.27 Flow cytometry size and fluorescence ranges of detection in nanometers and MESF, respectively | | | |
| 3.28 Brand and catalog number for ELISA kit used for each Apolipoprotein test (e.g. ApoB) | ApoA1: Mabtech, #3710-1HP-10<br>ApoB100: Mabtech, #3715-1HP-10 | | |
| 3.29 Concentration measured for each Apolipoprotein tested (mean µg/mL) | ApoA1: 0.102<br>ApoB100: 0.017 (mean of detectable samples; 92 samples were under the limit of detection and thus not included into this calculation) | | |
