## Supplementary material for "Comprehensive Phenotyping of Extracellular Vesicles in Blood of Healthy Humans – Insights into Cellular Origin and Biological Variability": MIBlood-EV serum

INTERNATIONAL SOCIETY for EXTRACELLULAR VESICLES

### MIBlood-EV

#### Standardized Reporting Tool for Blood EV Research (Human)

#### STUDY INFORMATION

| | |
|---|---|
| 1.0 Manuscript title | Comprehensive Phenotyping of Extracellular Vesicles in Blood of Healthy Humans – Insights into Cellular Origin and Biological Variability |
| 1.1 Corresponding author (Name and Email) | Metka Lenassi, |
| 1.2 Institution name | Institute of Biochemistry and Molecular Genetics, Faculty of Medicine, University of Ljubljana, Ljubljana, Slovenia; EU |
| 1.3 Time period of experiment (e.g. 2022-2024) | 7.2020 - 7.2021 |
| 1.4 Number of samples | 208 |
| 1.5 Cargo of interest | ☐ Vesicles ☑ Protein ☐ RNA ☐ DNA ☐ Other: lipoproteins |
| 1.6 Biospecimen type | ☐ Plasma ☑ Serum |
| 1.7 Biospecimen state | Frozen |
| 1.8 Source of frozen specimens | Prepared by the lab |
| 1.9 Years of collection (range) | 7.2020-11.2020 |

#### BLOOD COLLECTION AND PROCESSING

| | |
|---|---|
| 2.0 Patient fasting status | Non-Fasted |
| 2.1 Fasting length (e.g. hours/days) | / |
| 2.2 Anatomical access site | cubital vein |
| 2.3 Needle diameter (e.g. gauge) | 16G |
| 2.4 Blood volume collected (mL) | 5 |
| 2.5 Plasma anticoagulant | ☐ EDTA ☐ Citrate ☐ Heparin ☐ Other: / |
| 2.6 Serum tube type | No additive |
| 2.7 Serum clotting time (minutes) | 30 |
| 2.8 Time between collection and first centrifugation (range in hours) | 0.5-3.0 |
| 2.9 Transport temperature | Room temp (~22°C) |
| 2.10 Transport condition of tubes | Vertical |
| 2.11 Centrifuge brand and model | Eppendorf Centrifuge 5702 |
| 2.12 Bucket rotor type | Swing |
| 2.13 Number of centrifugation cycles | One |
| 2.14a 1st Centrifugation speed (RCF in x g) | 2000 |
| 2.14b 1st Centrifugation time (minutes) | 10 |
| 2.15 1st Rotor brake | Off |
| 2.16 1st Centrifugation temperature | Room temp (~22°C) |
| 2.17a 2nd Centrifugation speed (RCF in x g) | / |
| 2.17b 2nd Centrifugation time (minutes) | / |
| 2.18 2nd Rotor brake | |
| 2.19 2nd Centrifugation temperature | |
| 2.20 Additional processing steps (e.g. filtration) | / |
| 2.21 Storage tubes (brand, type, source, catalog number) | Greiner bio-one, Cryo.s Freezing Tubes, 2 ml, #122277 |
| 2.22 Storage temperature | -80°C |
| 2.23 Length of storage (range in years) | 1-1.5 |

#### PLASMA/SERUM QUALITY CONTROL

| | |
|---|---|
| 3.0 Number of freeze-thaw cycles (range) | 1 |
| 3.1 Thawing temperature | on ice |
| 3.2 Thawing duration (minutes) | 120 |

##### Hemolysis

| | |
|---|---|
| 3.3 Presence of hemolysis | Not tested |
| 3.4 Frequency of hemolyzed samples (e.g. <25%, 25-50%) | |
| 3.5 Method used | |
| 3.6 RBC count (Median, 95% CI, N) | |
| 3.7 RBC counter brand and type | |
| 3.8 Spectrophotometry hemoglobin concentration (mean g/L) | |
| 3.9 Spectrophotometer brand, model and wavelength measured (e.g. 414 nm) | |
| 3.10 Hemolized samples were discarded | |

#### Platelets

| | | |
|---|---|---|
| 3.11 Presence of platelets | Not Tested | 3.12 Method used (e.g. Flow Cytometry) |
| 3.13 Marker(s) used (e.g. CD61, CD41) | | |
| 3.14 Concentration (median, 95% CI, N) | | |
| 3.15 Platelet counter instrument brand, type and limit of detection (cells/L) | | |
| 3.16 Flow cytometer brand and type | | |
| 3.17 Flow cytometry size and fluorescence ranges of detection in nanometers and MESF, respectively | | |

#### Lipoproteins

| | | | |
|---|---|---|---|
| 3.18 Presence of lipoproteins | Yes | 3.19 Method used (WB, ELISA, FC) | Other: Nephelometry |
| 3.20 Spectrophotometry L-index | | | |
| 3.21 Spectrophotometer brand, model and wavelength measured (e.g. 700 nm) | | | |
| 3.22 WB Marker(s) used (e.g. Apo B) | | | |
| 3.23 Western blot images provided in manuscript? | | | |
| 3.24 Flow cytometry marker(s) used (e.g. ApoB) | | | |
| 3.25 Flow cytometry concentration (median, 95% CI, N) | | | |
| 3.26 Flow cytometer brand and type | | | |
| 3.27 Flow cytometry size and fluorescence ranges of detection in nanometers and MESF, respectively | Info about Nephelometry:<br><br>Instrument: Atellica NEPH 630, Siemens<br>Lipoproteins measured: ApoA1, ApoB | | |
| 3.28 Brand and catalog number for ELISA kit used for each Apolipoprotein test (e.g. ApoB) | | | |
| 3.29 Concentration measured for each Apolipoprotein tested (mean µg/mL) | | | |
